# BAP1/ASXL complex modulation regulates Epithelial-Mesenchymal Transition during trophoblast differentiation and invasion

**DOI:** 10.1101/2020.12.01.405902

**Authors:** Vicente Perez-Garcia, Pablo Lopez-Jimenez, Graham J Burton, Ashley Moffett, Margherita Y. Turco, Myriam Hemberger

## Abstract

Normal function of the placenta depends on the earliest developmental stages when trophoblast cells differentiate and invade into the endometrium to establish the definitive maternal-fetal interface. Previously, we identified the ubiquitously expressed tumour suppressor BRCA1-associated protein 1 (BAP1) as a central factor of a novel molecular node controlling early mouse placentation. However, functional insights into how BAP1 regulates trophoblast biology are still missing. Using CRISPR/Cas9 knockout and overexpression technology, here we demonstrate that the downregulation of BAP1 protein is essential to trigger epithelial-mesenchymal transition (EMT) during trophoblast differentiation associated with a gain of invasiveness. This function, which is conserved in mouse and humans, is dependent on the binding of BAP1 binding to Additional sex comb-like (ASXL1/2/3) proteins to form the Polycomb repressive deubiquitinase (PR-DUB) complex. Our results reveal that the physiological modulation of BAP1 determines the invasive properties of trophoblast, delineating a new role of the BAP1 PR-DUB complex in regulating early placentation.

## Introduction

The placenta is a complex organ essential for nutrient and oxygen exchange between the mother and the developing fetus. Normal placental function in humans depends on the earliest stages of development, when trophoblast cells proliferate and differentiate to form the villous tree and invade into the maternal decidua. Trophoblast invasion allows attachment of the placenta to the uterus and also mediates transformation of maternal spiral arteries, thereby ensuring an unimpeded blood flow into the intervillous space. This process is fundamentally important to secure an adequate supply of resources to the fetus. Several pregnancy complications such a miscarriage, pre-eclampsia, placenta accreta, and fetal growth restriction (FGR) are underpinned by a primary defect in trophoblast invasion (Brosens et al. 2011; Kaufmann, Black, and Huppertz 2003). Despite extensive research, the precise molecular mechanisms that regulate correct trophoblast differentiation and invasion remain poorly understood.

As part of the Deciphering the Mechanisms of Developmental Disorders (DMDD) programme, we found that placental malformations are highly prevalent in embryonic lethal mouse mutants (Perez-Garcia et al. 2018). This means that a significant number of genetic defects that lead to prenatal death may be due to abnormalities of placentation. In addition, we identified new molecular networks regulating early placentation. One of these molecular hubs is centred around the tumour suppressor BRCA1-associated protein1 (BAP1), a deubiquitinase enzyme (DUB) involved in the regulation of the cell cycle, cellular differentiation, cell death, gluconeogenesis and DNA damage response (Carbone et al. 2013). At the molecular level, BAP1 regulates a variety of cellular processes through its participation in several multiprotein complexes. Amongst others, BAP1 has been reported to interact with the BRCA1-BARD1 (BRCA1-associated RING domain I) complex, forkhead box 1 and 2 (FOXK1/2), host cell factor-1 (HCF-1), yin yang 1 (YY1), O-linked N-acetylglucosamine transferase (OGT), lysine (K)-specific demethylase 1B (KDM1B) and methyl-CpG binding domain protein 5 and 6 (MBD5 and MBD6) (Jensen et al. 1998; Misaghi et al. 2009; Baymaz et al. 2014; Dey et al. 2012; Nishikawa et al. 2009; Yu et al. 2010).

BAP1 also binds the epigenetic scaffolding proteins additional sex combs like-1/2/3 (ASXL1/2/3) to form the Polycomb Repressive-Deubiquitinase (PR-DUB) complex that exerts an essential tumour suppressor activity by regulating ubiquitination levels of histone H2A (H2AK119Ub) (Scheuermann et al. 2010). ASXL proteins are obligatory partners of BAP1 and this interaction is required for BAP1 activity (Campagne et al. 2019). Mutations and deletions in PR-DUB core subunits, *BAP1* and *ASXL*, are frequently associated with various malignancies (Carbone et al. 2013; Murali, Wiesner, and Scolyer 2013; Abdel-Wahab et al. 2012; Triviai et al. 2019; Micol et al. 2014).

*Bap1* knockout (KO) mouse conceptuses are embryonic lethal around midgestation (E9.5) and exhibit severe placental defects that likely contribute to the intrauterine demise (Perez-Garcia et al. 2018). Specifically, the placentas of *Bap1*-mutant conceptuses show defects in differentiation of the chorionic ectoderm into syncytiotrophoblast, a process required for the development of the labyrinth, the area of nutrient exchange in the mouse placenta. Although conditional reconstitution of gene function in the placenta but not the embryo did not rescue the intrauterine lethality, it substantially improved syncytiotrophoblast formation. Moreover, *Bap1*-mutant placentas show a striking overabundance of trophoblast giant cells (TGCs), the invasive trophoblast cell population. A similar bias towards the TGC differentiation pathway at the expense of the syncytiotrophoblast lineage was observed in *Bap1*-null mouse trophoblast stem cells (mTSCs), suggesting a critical role for BAP1 in regulating trophoblast biology (Perez-Garcia et al. 2018).

Recent reports have highlighted the possibility of co-evolution of shared pathways of invasion between trophoblast and cancer cells, in particular with regards to cell invasiveness and the capacity to breach basement membranes (Kshitiz et al. 2019; Costanzo et al. 2018). In both cases, the initial process is characterized by epithelial-mesenchymal transition (EMT) where epithelial cells lose their polarity and cell-cell adhesion and gain migratory and invasive properties of mesenchymal cells (Parast, Aeder, and Sutherland 2001; El-Hashash, Warburton, and Kimber 2010). Understanding the mechanisms by which BAP1 regulates trophoblast differentiation and invasion will be important not only to uncover new molecular pathways involved in placental development, but also to shed light into the signalling pathways altered in tumours where BAP1 is mutated.

Here we sought to determine the molecular mechanism by which BAP1 regulates trophoblast proliferation, differentiation and invasion. Using CRISPR/Cas9-generated *Bap1*^-/-^ mTSCs, we found that the deletion of *Bap1* does not affect their self-renewal capacity, but precociously promotes the EMT process even in stem cell conditions. Furthermore, we demonstrate that BAP1 down-regulation is required to trigger EMT; consequently, *Bap1* overexpression, mediated by CRISPR-Synergistic Activation Mediator (SAM)-induced activation of the endogenous locus, delayed trophoblast invasion. Analysis of the PR-DUB complex components *Asxl1* and *Asxl2* revealed that *Asxl1* was downregulated in parallel to BAP1, regulating its stability. In contrast, *Asxl2* showed the opposite expression pattern, with a concomitant increase as cells differentiated. Like *Bap1*^-/-^ mTSCs, both *Asxl1-* and *Asxl2*-mutant mTSCs failed to induce syncytiotrophoblast differentiation. Moreover, the functional characterization of BAP1 in the human placenta and human trophoblast stem cells (hTSCs) suggests that the role of BAP1 in regulating trophoblast differentiation is conserved in mouse and humans. Collectively, these data reveal a pivotal role of BAP1/ASXL complexes in triggering EMT as a requisite for trophoblast invasion and in regulating the finely-tuned balance of lineage-specific differentiation into the various trophoblast subtypes.

## Results

### Bap1 is highly expressed in undifferentiated trophoblast and down-regulated as cells enter the TGC lineage

To gain insight into the role of *Bap1* in trophoblast development, we first examined BAP1 expression in mTSCs. This unique stem cell type is derived from the trophectoderm of the blastocyst or from extra-embryonic ectoderm (ExE) of early post-implantation conceptuses. mTSCs retain the capacity to self-renew and to differentiate into all trophoblast subtypes under appropriate culture conditions (Tanaka et al. 1998). Immunofluorescence analysis of BAP1 in mTSCs showed strong nuclear staining (Figure 1A, 1B and 1C). We noticed that mTSC colonies containing areas of spontaneous differentiation, identified by decreased ESRRB stem cell marker expression, displayed a concomitant reduction in BAP1 staining intensity, suggesting that BAP1 is downregulated as trophoblast cells differentiate (Figure 1B). In line with these observations, differentiation of mTSCs *in vitro* revealed a significant reduction in BAP1 protein levels at days 3 and 6 compared to stem cell conditions, as shown by immunofluorescence staining and Western blot (WB) analysis (Figure 1C and 1D). The strongest downregulation was seen at day 6 when giant cells are the prevailing differentiated cell type (Murray, Sienerth, and Hemberger 2016; Perez-Garcia et al. 2018). However, *Bap1* mRNA levels did not significantly change across this differentiation time course, indicating that the functional regulation of BAP1 takes place at the post-transcriptional level (Figure 1E).

**Figure 1:**
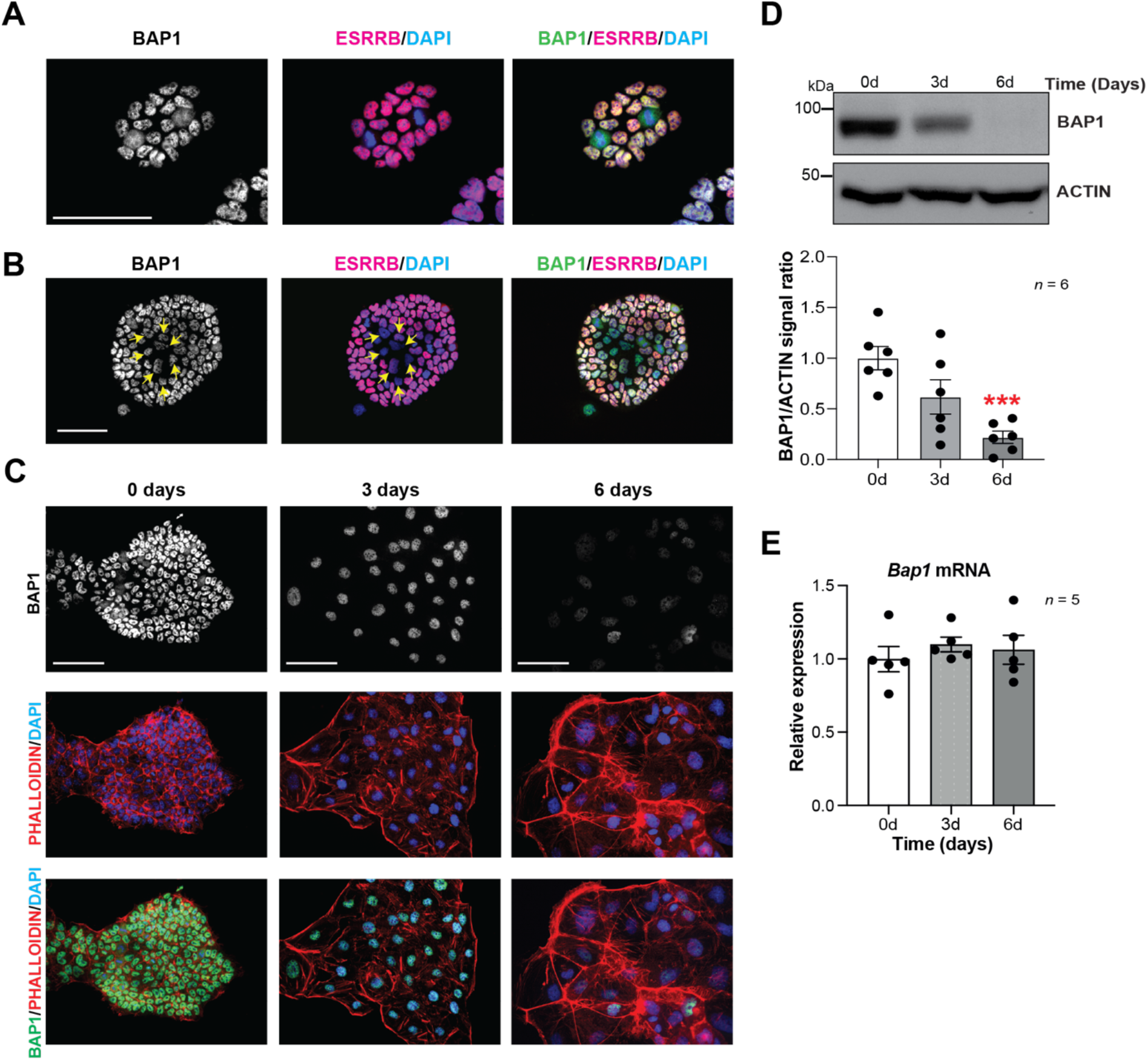
BAP1 protein levels are modulated during trophoblast differentiation. A, B) Immunofluorescence staining of mTSCs in the stem cell state for BAP1 and the stem cell marker ESRRB. The strong nuclear BAP1 staining observed in mTSCs is slightly reduced in partially differentiated, ESRRB-low cells (arrows). Scale bars: 100 µm. C) Immunofluorescence staining for BAP1 and F-Actin with phalloidin of mTSCs, mTSC differentiated for 3 and 6 days. BAP1 is downregulated as cells reorganize their cytoskeleton during trophoblast differentiation. Scale bars: 100 µm. D) Western blot for BAP1 on mTSCs in the stem cell state and upon 3 and 6 days of differentiation, confirming the downregulation of BAP1. Quantification of band intensities of six independent experiments is shown in the graph below. Data are normalized against ACTIN and relative to stem cell conditions (0d); Mean ± S.E.M.; ***p < 0.001 (one way-ANOVA with Dunnett’s multiple comparisons test). E) RT-qPCR analysis of Bap1 expression during a 6-day time course of mTSCs differentiation shows that Bap1 mRNA levels remain stable throughout the differentiation process. Expression is normalised to the Sdha and displayed relative to stem cell conditions (0d). Data are mean of five replicates ± S.E.M. (one way-ANOVA with Dunnett’s multiple comparisons test).

To further corroborate these results, we performed BAP1 immunostainings on mouse conceptuses at day (E) 6.5 of gestation, a time window when the ExE is actively proliferating and differentiating into the ectoplacental cone (EPC). While ExE will go on to develop predominantly into the labyrinth at later stages of development, EPC cells will give rise to the placental hormone-producing spongiotrophoblast layer and to invasive TGCs (Simmons, Fortier, and Cross 2007; Woods, Perez-Garcia, and Hemberger 2018). Immunofluorescence analysis revealed strong nuclear BAP1 staining in the embryo proper (epiblast, Epi) and in the ExE. However, differentiating EPC showed weak and diffuse staining (Figure 1-figure supplement 1A). In E9.5 placentae, BAP1 immunoreactivity was prominent in the developing labyrinth and spongiotrophoblast layer as well as in maternal decidual cells, but was markedly less pronounced in TGCs, again suggesting that BAP1 is specifically down-regulated as trophoblast cells differentiate into TGCs (Figure 1-figure supplement 1B).

Overall, these results indicate that BAP1 is highly expressed in undifferentiated trophoblast of the ExE *in vivo* and in mTSCs *in vitro*. BAP1 is downregulated at the protein level specifically as cells enter the TGC lineage, suggesting a potential function of BAP1 in regulating trophoblast differentiation and invasiveness, a key property of TGCs.

### BAP1 deletion does not impair the stem cell gene-regulatory network

The proliferative and self-renewal capacity of mTSCs depends on FGF and Tgfβ1/Activin A signalling pathways (Tanaka et al. 1998; Erlebacher, Price, and Glimcher 2004). To further explore the main growth factor signals involved in regulating BAP1 protein levels, mTSCs were subjected to 3 days of differentiation in the presence of either bFGF or conditioned medium (CM), which provides the main source of Tgfβ1/activin A in the complete TSC media. WB analysis showed that after 3 days of differentiation under standard differentiation conditions (Base medium) or in the presence of CM, BAP1 was markedly downregulated. However, the presence of bFGF alone maintained high BAP1 protein levels indicating that FGF signalling is dominant over Tgfβ1/Activin A in the positive regulation of BAP1 in stem cell conditions (Figure 2A).

**Figure 2:**
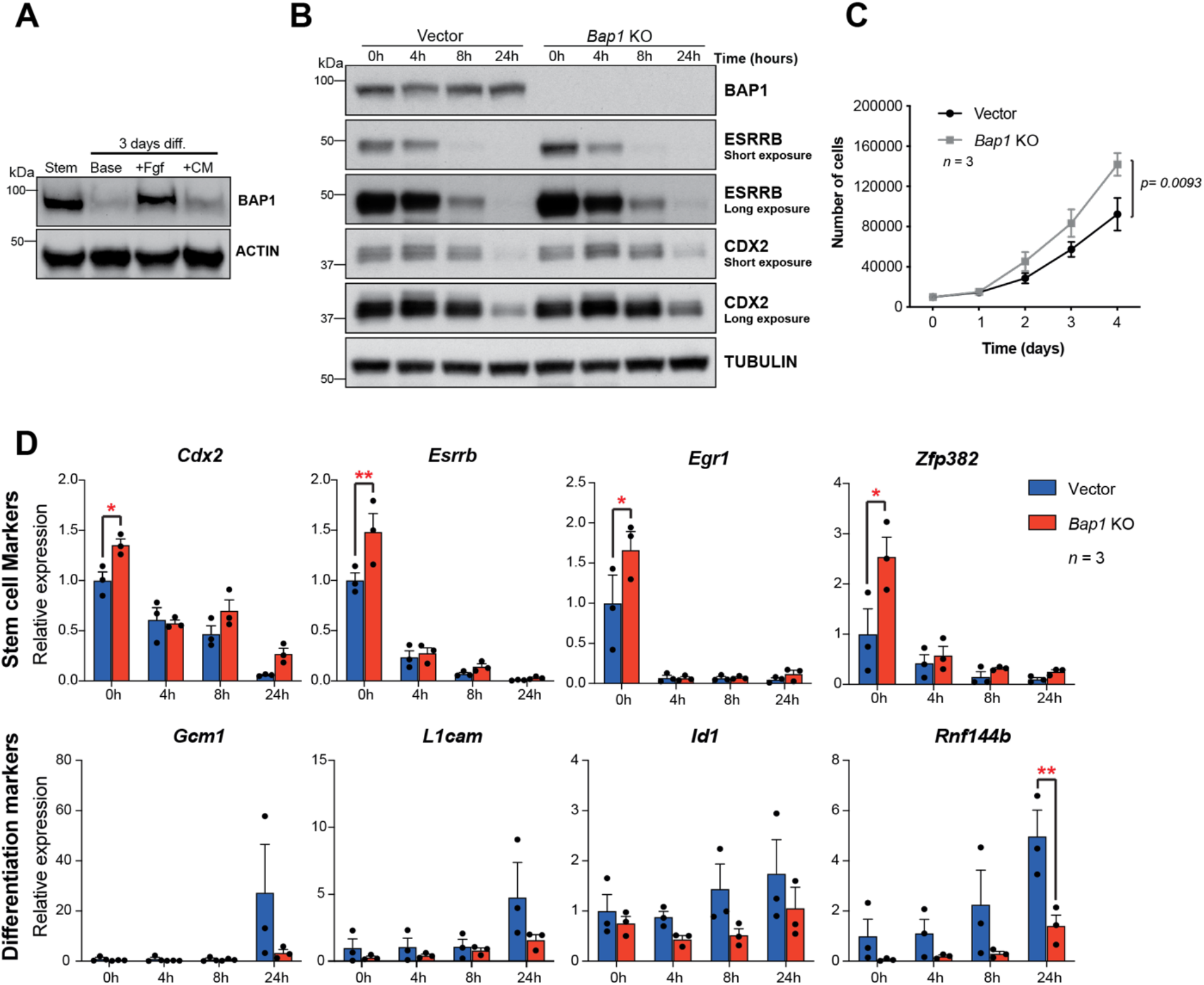
Bap1 ablation does not negatively affect stemness. A) Western blot analysis of mTSCs grown in stem cell conditions (Stem) and upon 3-day differentiation in standard Base medium (Base), or in base medium supplemented with FGF or conditioned medium (CM). B) Western blot analysis assessing the dynamic changes in the stem cell markers CDX2 and ESRRB across a short-term differentiation time course in vector control compared to Bap1-mutant mTSCs (stem cell conditions = 0h, and differentiation at 4h, 8h and 24h). Blots are representative of two independent replicates. C) Proliferation assay of control and Bap1^-/-^ mTSCs over 4 consecutive days. Bap1^-/-^ mTSCs exhibit a significant increase in the proliferation rate compared to vector control cells (mean ± S.E.M.; n = 3). p = 0.0093; two-way ANOVA with Holm-Sidak’s multiple comparisons test. D) RT-qPCR analysis of control and Bap1^-/-^ mTSCs for stem cell and early differentiation marker genes. Stem cell markers are increased and differentiation delayed in Bap1-mutant mTSCs. Data are normalised to Sdha and displayed as mean of 3 biological (clones) replicates ± S.E.M.; *p < 0.05, **p < 0.01 (two-way ANOVA with Sidak’s multiple comparisons test).

This raises the question whether the absence of BAP1 affects stem cell fate. The transcription factors CDX2 and ESRRB represent primary targets and direct mediators of FGF signalling in mTSCs (Latos et al. 2015b) that are essential to keep mTSCs in a highly proliferative, undifferentiated state. Both factors are rapidly downregulated upon trophoblast differentiation (Latos et al. 2015a; Luo et al. 1997; Strumpf et al. 2005). We assessed CDX2 and ESRRB protein levels in *Bap1*^*-*^*/*^*-*^ mTSCs (Perez-Garcia et al. 2018) compared to (empty vector) control cells across 24 hours of differentiation (0h=stem cell conditions; 4h, 8h, 24h = hours upon differentiation). The absence of BAP1 resulted in increased ESRRB and CDX2 protein levels, and increased proliferation rates in stem cell conditions (Figure 2B and 2C). The higher residual expression of ESRRB and CDX2 proteins detected after 24h of differentiation may indicate a potential delay in mTSC differentiation (Figure 2B). Indeed, analysis of the mRNA expression dynamics of the trophoblast stem cell markers *Cdx2, Esrrb, Egr1* and *Zpf382* indicated that *Bap1*-mutant mTSCs differentiated more slowly than the control counterparts during the initial 24 hours of differentiation (Figure 2D and Figure 2-figure supplement 1A). In line with these results, we also observed that the upregulation of early mTSC differentiation markers such as *Gcm1, L1cam, Id1* and *Rnf44b* was delayed in *Bap1*^-/-^ mTSCs compared to control cells (Figure 2D and Figure 2-figure supplement 1A).

### Bap1^-/-^ mTSCs undergo Epithelial-Mesenchymal Transition (EMT)

The appearance of *Bap1*^-/-^ TSCs under phase contrast reveals a phase bright, refractile and loosely associated morphology with poor cell-cell contacts. This is in contrast to the colonies of vector control cells, suggesting that they may have undergone EMT-like transition (Figure 3A), known to occur when trophoblast differentiates towards the invasive TGC lineage (Sutherland 2003). The morphology of *Bap1*^-/-^ mTSC colonies led us to hypothesize that BAP1 affects EMT in trophoblast. To investigate this, we studied the global expression profile of *Bap1*-mutant TSCs compared to control cells in stem cell conditions (0d) and upon differentiation (3d). Unbiased clustering and principal component analysis clearly showed that the differentially expressed genes (DEG) were determined by the absence of BAP1 and by the day of differentiation (Figure 3B and Figure 3-figure supplement 1A). Gene Ontology analysis revealed an enrichment of genes involved in regulation of extracellular matrix, cell junction and cell adhesion at both time points analysed, concordant with the marked change in morphology of *Bap1*^-/-^ mTSCs (Figure 3C, Figure 3-figure supplement 1B and Supplementary Table1 and 2). In line with these results, *Bap1*^-/-^ mTSCs exhibited a significant decrease in cell adhesion on tissue culture plastic (Figure 3D), which was even more obvious when *Bap1*^-/-^ mTSCs were grown in 3D organoid-like trophospheres (Figure 3E) (Rai and Cross 2015b).

**Figure 3:**
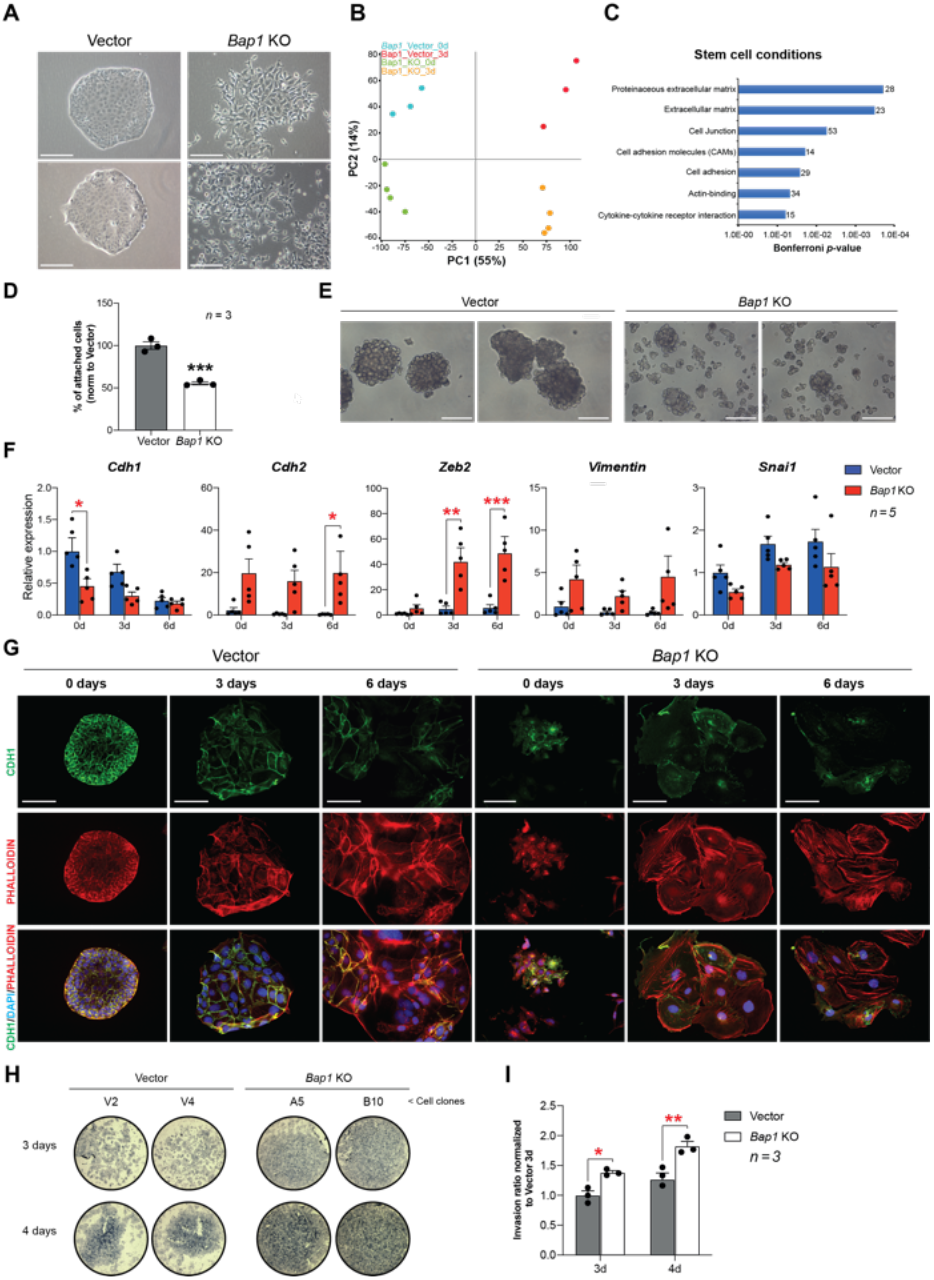
Bap1-deficiency promotes epithelial-mesenchymal transition (EMT). A) Colony morphology of wild type (vector) and Bap1-mutant mTSCs. Bap1^-/-^ mTSCs show a fibroblast-like morphology with loss of cell-cell attachment compared to vector control mTSCs. Scale bar: 100 µm. B) Principal component analysis of global transcriptomes of independent vector control (n=3) and Bap1 KO (n=4) clones grown in stem cells conditions (0d) and after three days of differentiation (3d). C) Gene ontology analyses of genes differentially expressed between vector and Bap1-mutant mTSCs in stem cell conditions. D) Cell adhesion assay showing that Bap1-mutant mTSCs are less well attached to cell culture plastic compared to vector control cells. Data are mean of three independent replicates with 3 biological replicates (=independent clones) per experiment. ***p < 0.001 (Student’s t-test). E) Morphology of 3D-trophospheres after 8 days of differentiation in low attachment conditions. Scale bar: 200 µm. F) RT-qPCR analysis of EMT marker expression during a 6-day differentiation time course. Data are normalised to Sdha and displayed relative to vector 0d. Data are mean of 5 biological (= independent clones) replicates ± S.E.M.; *p < 0.05, **p < 0.01, ***p < 0.001 (two-way ANOVA with Sidak’s multiple comparisons test). G) Immunofluorescence analysis for CDH1 and F-Actin (phalloidin) of vector control and Bap1-mutant mTSCs over 6 days of differentiation. Lack of BAP1 reduces cell-cell junctions (CDH1 staining) with a profound reorganization of the cytoskeleton (increased actin stress fibres). Data are representative of 5 independent vector control and Bap1 KO clones each. Scale bar: 100 µm. H) Transwell invasion assay of vector control (V2, V4) and Bap1-mutant (clones A5, B10) mTSCs after 3 and 4 days of differentiation. Photographs of invasion filters show haematoxylin-stained cells that reached the bottom side of the filter after removal of the reconstituted basement membrane matrix (Matrigel). I) Quantification of invaded cells, measured by the colour intensity, normalized to 3d controls. Data are mean of three independent replicates (3 biological clones in each replicate) ± S.E.M.; *p < 0.05, **p < 0.01** (two-way ANOVA with Sidak’s multiple comparisons test).

At the molecular level, the reduction in cell adhesion was marked by a significant downregulation of E-Cadherin (*Cdh1*), an epithelial hallmark, in *Bap1*^-/-^ compared to vector control cells (Figure 3-figure supplement 1D and Supplementary Table 1 and 2). Stringent calling (DESeq2 and Intensity difference analysis) of DEG revealed that several genes involved in the stabilization of cell-cell contacts and epithelial integrity (*Claudin 4* (*Cldn4), Claudin 7* (*Cldn7), Desmoplakin* (*Dsp)*, and *Serpine1)* were down-regulated in *Bap1*-mutant mTSCs (Figure 3-figure supplement 1C and Supplementary Table 1). Concomitant with the downregulation of epithelial markers like *Cdh1*, mesenchymal markers including N-Cadherin (*Cdh2), Zeb2* and *Vimentin (Vim)* were upregulated in the absence of *Bap1* (Figure 3F). These data indicate that *Bap1*-null mTSCs display a pronounced and precocious EMT phenotype.

TGC formation is characterized by cytoskeletal rearrangement, exit from the cell cycle, DNA endoreduplication and production of trophoblast-specific proteins such as placental prolactins. Thus, undifferentiated trophoblast cells exhibit little organised actin and few peripheral focal complexes, whereas TGCs show a highly organised cytoskeleton containing prominent actin stress fibres linked to gain in motility and invasiveness (Parast, Aeder, and Sutherland 2001; El-Hashash, Warburton, and Kimber 2010). As expected from the mRNA expression analysis, *Bap1*^-/-^ mTSCs showed a loss of membrane-associated CDH1 staining and disorganized cytoskeleton in stem cell conditions, with increased numbers of actin stress fibres upon differentiation, suggesting a more TGC-like and invasive phenotype compared to wild-type (vector) cells (Figure 3G). Indeed, *Bap1*^-/-^ TSCs were also more invasive through extracellular basement membrane (Matrigel) compared to vector control mTSCs in Transwell invasion experiments (Figure 3H and 3I). In line with these results, the differentially expressed genes in *Bap1*^-/-^ mTSCs showed significant overlap with the gene expression signatures of tissues prone to form tumours such as *Bap1*^-/-^ melanocytes and mesothelial cells (He et al. 2019) (Figure 3-figure supplement 1E). Altogether, these results indicate that the lack of *Bap1* triggers an EMT in mTSCs that recapitulates critical aspects of early malignant transformation.

### Levels of Bap1 are critical to trigger EMT during trophoblast differentiation

In order to confirm that BAP1 is one of the main drivers of EMT during trophoblast differentiation, we overexpressed BAP1 using the CRISPR/gRNA-directed Synergistic Activation Mediator (SAM) technology (Konermann et al. 2015). One out of three single guide RNAs (sgRNAs) tested induced robust upregulation of *Bap1* mRNA and BAP1 protein levels compared to mTSCs transduced with non-targeting sgRNA (NT-sgRNA) (Figure 4A, 4B and Figure 4-figure supplement 1A). The morphology of *Bap1*-overexpressing mTSCs was characterized by tight epithelial colonies compared to NT-sgRNA control mTSCs (Figure 4C). To gain insight into the global transcriptional response to *Bap1* overexpression, we performed RNA-seq on mTSCs grown in stem cell conditions (0d) and after 3 days of differentiation (3d). PCA analysis showed that, besides the growth conditions, samples clearly cluster by the levels of BAP1 within the cells (Figure 4-figure supplement 1B and 1C). A stringent assessment of the deregulated genes (DESeq2 and intensity difference filter) revealed that, in addition to *Bap1*, a cohort of 80 genes were significantly deregulated with a robust upregulation of genes involved in cell junction biology and maintenance of epithelial integrity – such as *Plakophilin 2* (*Pkp2*), *Keratin-7/8/19* (*Krt7/8/19*), *Desmoplakin* (*Dsp*) and *Cingulin* (*Cgn*) (Figure 4D and Supplementary Table 3 and 4). In line with these observations, Gene Ontology analysis revealed an overrepresentation of extracellular matrix and cell adhesion molecules, suggesting an increase in epithelial features of BAP1-overexpressing cells compared to control cells (Figure 4E and Supplementary Table 3 and 4).

**Figure 4:**
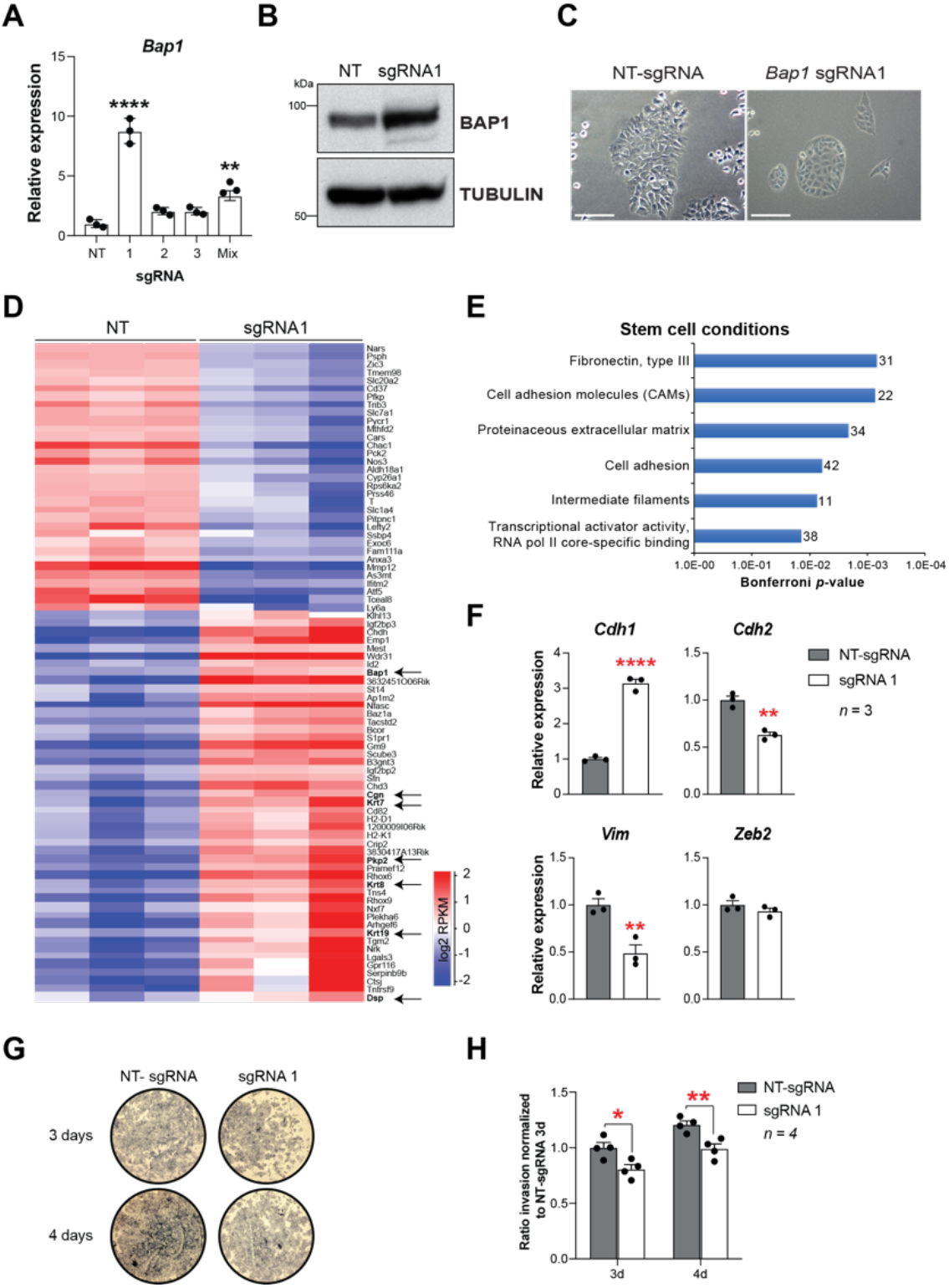
Bap1 overexpression enhances epithelial features and reduces invasiveness. A) RT-qPCR analysis to determine the overexpression of Bap1 in mTSCs induced by the transduction of each single guide RNAs (sgRNAs) or in combination (Mix) compared to non-targeting sgRNA (NT). The strongest Bap1 upregulation was observed following transduction with sgRNA1. Data are normalised to Sdha and are displayed as mean of three replicates ± S.E.M.; **p < 0.01, ****p < 0.0001 (one way-ANOVA with Dunnett’s multiple comparisons test). B) Confirmation of BAP1 overexpression assessed by Western blot. Tubulin was used as loading control. C) Colony morphology of non-targeting (NT) sgRNA and sgRNA1-transduced mTSCs. Overexpression of BAP1 in sgRNA1 mTSCs increases epithelioid features of the cell colonies. D) Heatmap of differentially expressed genes in mTSCs transduced with non-targeting (NT) sgRNA compared sgRNA1. Arrows point to Bap1 itself and to genes associated with the reinforcement of epithelial integrity. Three independent biological replicates per genotype were sequenced. E) Gene ontology analyses of genes differentially expressed between sgRNA1 and NT-sgRNA mTSCs grown in stem cell conditions. F) RT-qPCR analysis of epithelial and mesenchymal markers in NT-sgRNA control cells compared to sgRNA1 Bap1-overexpressing mTSCs. Data are normalised to Sdha and are displayed as mean of three replicates ± S.E.M.; **p < 0.01, ****p < 0.0001 (Student’s t-test). G) Transwell invasion assays of control and Bap1 overexpressing mTSCs. Representative images are shown. H) Quantification of invaded cells, measured by the colour intensity, normalized to 3d controls. Data are mean of four independent replicates ±S.E.M.; *p < 0.05, **p < 0.01 (two-way ANOVA with Sidak’s multiple comparisons test).

To corroborate these results, we validated several EMT markers by RT-qPCR and confirmed that the overexpression of *Bap1* induced a significant upregulation of *E-cadherin* (*Cdh1*) with concomitant downregulation of *N-cadherin* (*Cdh2*) and *Vimentin* (*Vim*) (Figure 4F). These results demonstrated that precise levels of BAP1 regulate mTSCs morphology, and that modulation of BAP1 levels affects the extent and speed at which trophoblast cells undergo EMT (Figure 4-figure supplement 1D and Supplementary Table 3 and 4). In line with the re-acquisition of epithelial properties, *Bap1*-overexpressing mTSCs showed lower invasive capacity through Matrigel compared to NT-sgRNA control cells (Figure 4G and 4H). Altogether these results indicate that the downregulation of BAP1 is critical for triggering EMT and invasiveness of trophoblast cells.

### BAP1 and ASXL1/2 complexes are co-regulated during trophoblast differentiation

Interaction of ASXL proteins with BAP1 promotes its stability and enzymatic activity (Campagne et al. 2019). In order to investigate the role of the BAP1-ASXL complex in regulating trophoblast biology, we first examined gene expression of ASXL family members *Asxl1* and *Asxl2* over a 6-day differentiation time course. RT-qPCR and WB analysis showed that ASXL1 was highly expressed in TSCs under stem cell conditions and strongly downregulated during trophoblast differentiation in parallel to BAP1 protein levels. ASXL2 expression displayed the opposite trend with maximal levels in differentiated trophoblast (Figure 5A and 5B). This expression pattern was further validated by immunofluorescence (Figure 5-figure supplement 1A and 1B). To gain insight into the specific roles of ASXL1 and ASXL2 we generated KO mTSCs using CRISPR/Cas9 technology (Figure 5-figure supplement 1C). The deletion of *Asxl1* or *Asxl2* did not affect stemness. However, under differentiation conditions, *Asxl1*^-/-^ and *Asxl2*^-/-^ mTSCs failed to upregulate markers of syncytiotrophoblast, whereas the differentiation towards TGCs was promoted (Figure 5C, 5D and Figure 5-figure supplement 1D). This defect phenocopied the syncytiotrophoblast differentiation defect we had previously reported for *Bap1*-mutant cells (Perez-Garcia 2018). Moreover, the absence of *Asxl1* and *Asxl2* induced an upregulation of EMT markers such *Cdh2, Vim, Zeb1* and *Zeb2* suggesting that ASXL1 and ASXL2 together with BAP1 contribute to modulating EMT during trophoblast differentiation (Figure 5-figure supplement 1D).

**Figure 5:**
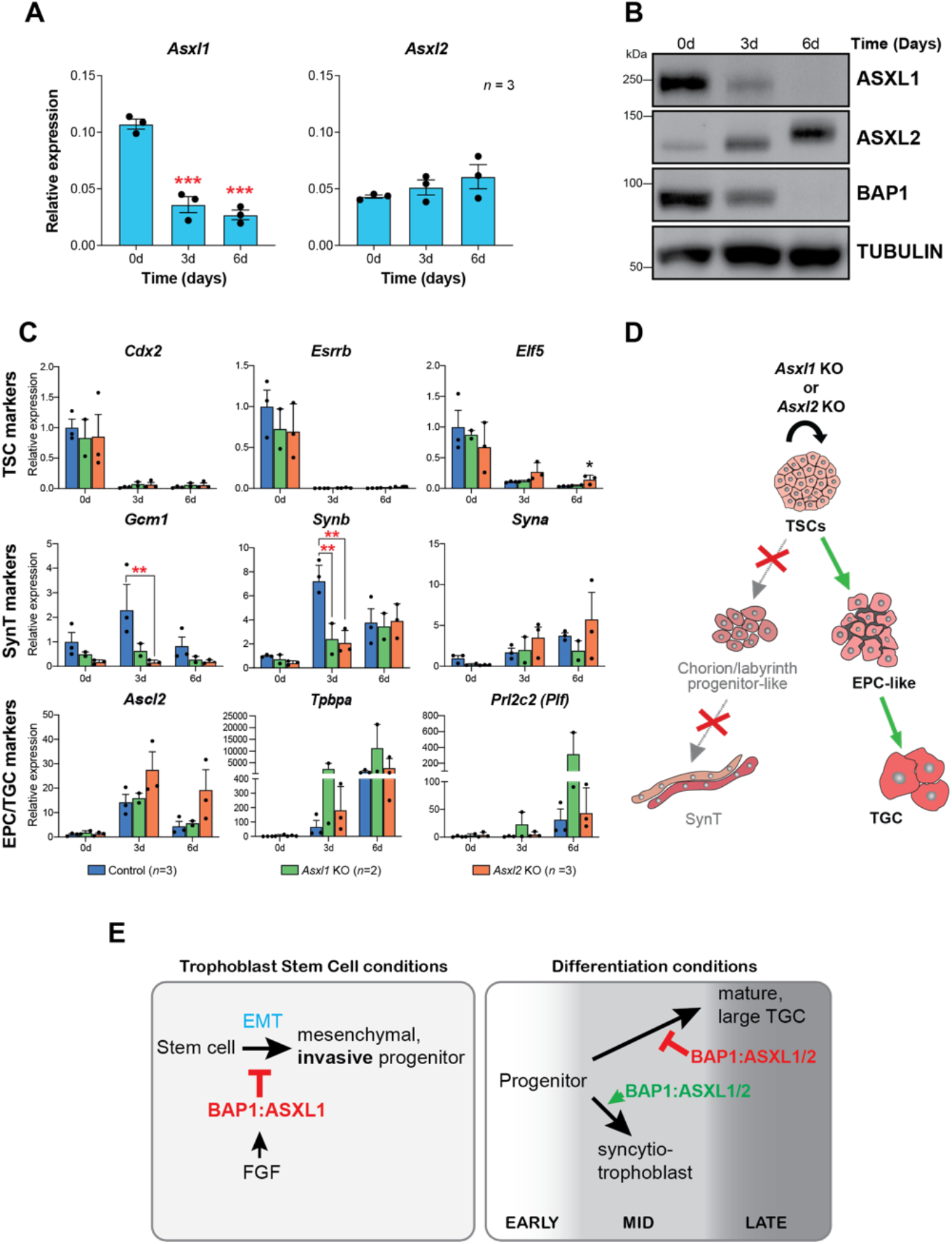
BAP1 and ASXL1/2 complexes are co-regulated during trophoblast differentiation. A) RT-qPCR analysis of Asxl1 and Asxl2 expression during a 6-day mTSC differentiation time course. Data are normalised to Sdha and are displayed as mean of 3 replicates ± S.E.M.; ***p < 0.001 (one way-ANOVA with Dunnett’s multiple comparisons test). B) Western blot analysis of ASXL1 and ASXL2 protein levels in mTSCs differentiating over 6 days. Blots shown are representative of 3 independent replicates. C, D) Analysis of Asxl1^-/-^ and Asxl2^-/-^ mTSCs grown in self-renewal conditions (0d) or after differentiation for 3 and 6 days assessed by RT-qPCR. Data are mean ± S.E.M. of n = 3 (control, scramble), n = 2 (Asxl1 KO) and n = 3 (Asxl2 KO) individual clones as independent replicates. **p < 0.01 (two-way ANOVA with Sidak’s multiple comparisons test). D) Schematic diagram of the differentiation defects observed in Asxl1^-/-^ and Asxl2^-/-^ mTSCs. E) Suggested model for the role of PR-DUB BAP1/ASXL complexes in regulating trophoblast biology. In stem cell conditions, the presence of FGF stimulates the stability of the BAP1/ASXL1 complex, which maintains the epithelial integrity of the mTSCs preventing EMT. Upon differentiation (FGF withdrawal), BAP1/ASXL1 is reduced with a concomitant upregulation of ASXL2 protein levels as differentiation progresses. Notably, from early to mid-differentiation both BAP1:ASXL1 and BAP1:ASXL2 complexes coexist and are essential to trigger syncytiotrophoblast differentiation.

### BAP1 PR-DUB complex is also regulated during human trophoblast differentiation

TGCs represent the invasive trophoblast cell type in mice whereas in humans this function is exerted by extravillous trophoblast (EVT). As in mouse, the gain of invasive properties is accompanied by an EMT process (DaSilva-Arnold et al. 2015; J et al. 2016; Vicovac and Aplin 1996). Polycomb complexes, including the BAP1 PR-DUB, are well conserved throughout evolution (Chittock et al. 2017) leading to the question whether BAP1 also functions to regulate trophoblast differentiation and invasion during human placentation.

To determine the dynamics of BAP1 expression in human trophoblast we isolated RNA from placental villi across gestation, and performed RT-qPCRs to analyse *BAP1* expression. Despite some variability, *BAP1* mRNA levels increased over pregnancy (Figure 6A). The expression of BAP1 was also analysed in human trophoblast stem cells (hTSCs) and choriocarcinoma cell lines. Interestingly, among the placental choriocarcinoma cell lines, the most invasive cell line JEG-3 (Grummer et al. 1994) showed lowest BAP1 expression levels compared to JAR and BeWo cell lines, suggesting that BAP1 may also play a role in regulating human trophoblast invasion (Figure 6B). In first-trimester placentae, strong expression of BAP1 was observed in villous cytotrophoblast (VCT) and at the base of cytotrophoblast cell columns (CCC) compared to the very low signal in syncytiotrophoblast (SCT) (Figure 6C). Of note, BAP1 staining became markedly weaker and more diffuse along the distal aspects of the CCC as cells undergo EMT and differentiate towards invasive EVT (Figure 6C). High expression of integrin alpha-5 (ITGA5), a marker of EVT, correlated with decreased staining intensity of BAP1, suggesting that BAP1 was downregulated during EVT differentiation (Figure 6-figure supplement 1A).

**Figure 6:**
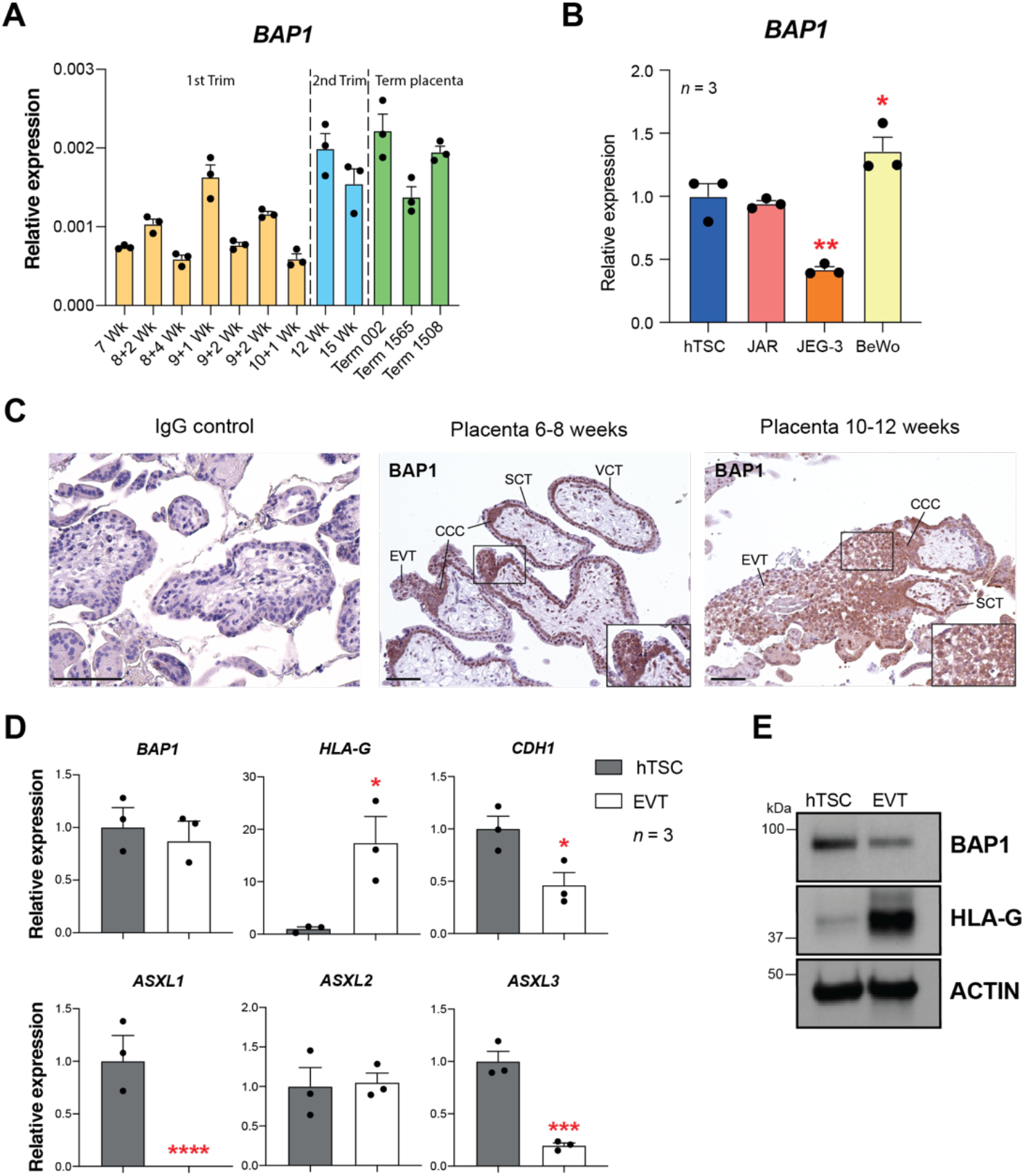
BAP1 PR-DUB modulation is also observed in human placentation. A) RT–qPCR analysis of BAP1 expression on human placental villous samples ranging from 7 weeks of gestation to term. Three independent term placental samples were investigated. An overall increase of BAP1 expression was observed over gestation. Expression is normalised to YWHAZ housekeeping gene. Data are mean of 3 replicates ± S.E.M. B) RT-qPCR analysis of BAP1 expression in human trophoblast stem cells (hTSCs) and the choriocarcinoma cell lines JAR, JEG-3 and BeWo. Expression is normalised to GAPDH. Data are mean of 3 replicates ± S.E.M.; *p< 0.05 **p < 0.01 (one way-ANOVA with Dunnett’s multiple comparisons test). C) Immunohistochemistry for BAP1 on early first trimester placenta (6-8 weeks g.a) and late first trimester (10-12 weeks g.a.). BAP1 staining is strong in proliferative villous cytotrophoblast (VCT) and cytotrophoblast cell columns (CCC) compared to syncytiotrophoblast (SCT). Notably, invasive extravillous trophoblast (EVT) shows a diffuse and weak staining as cells undergo EMT. Scale bars, 100µm. D) RT-qPCR analysis of BAP1, HLA-G, CDH1 and ASXL1-3 gene expression on hTSCs and in vitro-differentiated EVT cells after 8 days of differentiation. Expression is normalised to GAPDH. Data are mean of 3 independent replicates ± S.E.M.; *p< 0.05, ***p < 0.001, ****p < 0.0001 (Student’s t-test). E) Western blot analysis of BAP1 protein levels in EVT compared to hTSCs. As in the mouse, BAP1 is strongly reduced during trophoblast differentiation towards the invasive EVT lineage. Blots shown are representative of 3 independent replicates.

To further corroborate these results we differentiated hTSCs towards EVT for 8 days (Okae et al. 2018) and examined the expression of BAP1/ASXL complex components by RT-qPCR. We confirmed successful EVT differentiation by *HLA-G* expression and concomitant downregulation of *CDH1*. Although *BAP1* mRNA expression levels remain unchanged, protein levels declined markedly upon EVT differentiation (Figure 6D and 6E), in line with the post-transcriptional regulation of BAP we had observed in the mouse (Figure 1D and 1E). In addition to *ASXL1* and *ASXL2*, the *ASXL3* isoform was also expressed in hTSCs, and together with ASXL1, was significantly downregulated upon EVT differentiation (Figure 6D). Taken together, these results indicate that the molecular mechanism by which the BAP1/ASXL complexes regulate trophoblast differentiation and invasion may be conserved in human and in mice.

## Discussion

The similarities between trophoblast and tumour cells have long been recognised, in particular with respect to their invasive properties (Costanzo et al. 2018; Ferretti et al. 2007). BAP1, a tumour suppressor frequently mutated in human cancers, is ubiquitously expressed and inactivation or deletion of this gene results in metastasis (Carbone et al. 2013). During murine development, embryos deficient for *Bap1* die during the stage of early organogenesis (E9.5) with severe placental dysmorphologies. Although a central role for BAP1 during early placentation was suggested (Perez-Garcia et al. 2018), its function in regulating trophoblast development has not been explored. In this study, we show that BAP1 is highly expressed in both mTSCs and hTSCs, and that its downregulation triggers EMT and promotes trophoblast invasiveness. We also find that BAP1 protein levels are tightly coordinated with the expression of the ASXL proteins, indicating that modulation of the PR-DUB complex is required for proper trophoblast differentiation. To decipher the mechanism by which BAP1 and ASXL may regulate trophoblast self-renewal and differentiation, we deleted and overexpressed these factors in mTSC using CRISPR/Cas9-KO and CRISPR/Cas9-SAM activation technology- the first of its kind in mTSCs to date. We find that BAP1 regulates many facets of trophoblast biology. Firstly, in stem cell conditions, *Bap1* ablation triggers an overt EMT phenotype associated with increased cellular invasiveness, and, at the same time, enhances proliferation and expression of stem cell markers. This suggests that deficiency of *Bap1* uncouples the normal loss of proliferation from differentiation, reminiscent of malignant transformation. Indeed, the upregulation of stem cells markers upon functional BAP1 depletion is seen in human uveal melanoma and renal cell carcinoma, and is associated with aggressive cancer behaviour and poor patient outcome (Matatall et al. 2013; Pena-Llopis et al. 2012; Harbour et al. 2010). In line with these data, overexpression of BAP1 induces the converse phenotype in mTSCs with reinforcement of epithelial features and reduced invasiveness. Therefore, we propose that BAP1 modulation is one of the main drivers triggering the EMT and invasion processes in trophoblast. In line with this view, BAP1 mutations in human liver organoids result in loss of cell polarity, epithelial disruption and increased cell motility, features observed during the initial steps of EMT (Kalluri and Weinberg 2009; Das et al. 2019).

In the absence of an FGF signal, *Bap1*-deficiency promotes trophoblast differentiation towards a TGC phenotype, and represses syncytiotrophoblast formation. This is shown by the precocious cytoskeletal rearrangements we describe in mutant mTSCs, as well as the profound over-abundance of terminally differentiated, extremely large TGCs in the KO placentae (Perez-Garcia et al. 2018). These dual roles of BAP1 depending on the FGF signalling environment may indeed be explained by the differential regulation of its PR-DUB components, ASXL1 and ASXL2. BAP1 and ASXL proteins form mutually exclusive complexes of the PR-DUB tumour suppressor, which maintains transcriptional silencing of Polycomb target genes. Moreover, because ASXL proteins can affect the stability of BAP1, the high discrepancy between mRNA and protein levels in TSCs can be explained (Scheuermann et al. 2010; Daou et al. 2015). Our data suggest that the BAP1:ASXL1 complex is predominant in TSCs and plays an important role in preventing mTSCs from undergoing EMT while in a proliferative stem cell state. With the onset of trophoblast differentiation and the concomitant upregulation of ASXL2, both BAP1:ASXL1 and BAP1:ASXL2 complexes will coexist. Both these complexes are important to promote syncytiotrophoblast differentiation; in the absence of either *Bap1, Asxl1* or *Asxl2*, syncytiotrophoblast differentiation is abrogated, and TGC differentiation dominates (Figure 5E). This is in keeping with the finding that overexpression of *Asxl2* induces cellular senescence in other systems (Huether et al. 2014; Micol et al. 2014; Daou et al. 2015).

We previously found a strong correlation between cardiovascular and brain defects in embryos with abnormal placentation (Woods, Perez-Garcia, and Hemberger 2018). Since *Asxl1* and *Asxl2* mutants have also been reported to exhibit cardiovascular and brain developmental defects (Baskind et al. 2009; Wang et al. 2014), it is tempting to speculate that they may be due, in part, to a placental defect.

Finally, our data suggest that BAP1:ASXL1/2 regulate trophoblast differentiation and invasiveness in other species. In humans as in mice, BAP1 protein levels are downregulated during differentiation towards the invasive EVT lineage in coordination with ASXL gene expression. We also observed that in addition to *ASXL1* downregulation, *ASXL3* expression was also modulated during EVT differentiation. *De novo* mutations of *ASXL1/2/3* genes are associated with severe fetal growth restriction, preterm birth and defects in the development of the heart-brain axis (Srivastava et al. 2016), i.e. defects strongly linked to abnormal placentation. Our work suggests a direct link of these mutations to abnormal trophoblast development through the various functions of PR-DUB in regulating the unique properties of trophoblast cells. Gaining detailed insights into the molecular networks regulating this BAP1-ASXL modulation during early placentation will help not only to shed light onto the major unexplained pregnancy disorders, but also to open up new avenues into investigations of tumours where PR-DUB is mutated.

## Methods

### Cell culture and generation of mutant TSC lines

The wild-type TS-Rs26 TSC line (a kind gift of the Rossant lab, Toronto, Canada) and mutant TSC lines were grown as previously described (Tanaka et al. 1998). Briefly, mTSCs were cultured in standard mTSCs conditions: 20% foetal bovine serum (FBS) (ThermoFisher Scientific 10270106), 1 mM sodium pyruvate (ThermoFisher Scientific 11360-039), 1X Anti-mycotic/Antibiotic (ThermoFisher Scientific 15240-062), 50 µM 2-mercaptoethanol (Gibco 31350), 37.5 ng/ml bFGF (Cambridge Stem Cell Institute), and 1 µg/ml heparin in RPMI 1640 with L-Glutamine (ThermoFisher Scientific 21875-034), with 70% of the medium pre-conditioned on mouse embryonic fibroblasts (CM). The medium was changed every two days, and cells passaged before reaching confluency. Trypsinisation (0.25% Trypsin/EDTA) was carried out at 37 °C for about 5 min. Differentiation medium consisted of unconditioned TSC medium without bFGF and heparin.

*Bap1* KO mTSC clones were generated in our laboratory and published before (Perez-Garcia et al. Nature 2018). *Bap1* was overexpressed in mTSCs by using CRISPR/Cas9 Synergistic Activation Mediator system (Konermann et al. 2015). In brief, SAM mTSCs were generated by lentiviral transduction of lenti dCas9-VP64-Blast (Addgene 61425) and lenti MS2-p65-HSF1-Hygro (Addgene 61426) into TS-Rs26 mTSCs, followed by antibiotic selection. Then, to generate SAM *Bap1* mTSCs, three *Bap1*-gRNA targeting the 180 bp region upstream of the *Bap1* TSS and one non-targeting-gRNA (Supplementary Table 5) (Joung et al. 2019) were selected and synthesized (Sigma). Each oligo was annealed and cloned into the sgRNA (MS2)-puro plasmid (Addgene 73795) by a Golden Gate reaction using BsmBI enzyme (Thermo Fisher Scientific, ER0451) and T7 ligase (NEB, M0318S). The new gRNA constructs were packaged into lentiviral particles and transduced into SAM TSCs by direct supplementation of the lentivirus for 24h. After 48h, SAM *Bap1* overexpressing cells were selected by adding 1µg/1ml puromycin for 7 days.

CRISPR/Cas9-mediated *Asxl1*- and *Asxl2*-mutant mTSCs were generated as in (Lopez-Tello et al. 2019). Briefly, non-targeting gRNA (control) and gRNAs (Supplementary Table 5) that result in frameshift mutations were designed using the CRISPR.mit.edu design software and cloned into the Cas9.2A.EGFP plasmid (Plasmid #48138 Addgene). Transfection of gRNA Cas9.2A.EGFP constructs was carried out with Lipofectamine 2000 (ThermoFisher Scientific 11668019) reagent according to the manufacturer’s protocol. KO clones were confirmed by genotyping using primers spanning the deleted exon, and by RT-qPCR with primers within, and downstream of, the deleted exon, as shown (Figure 5-figure supplement 1C).

BTS5 blastocyst-derived hTSCs were donated by Prof Takahiro Arima and cultured as in (Okae et al. 2018). Briefly, hTSC were grown in 5 mg/ml Col IV coated 6-well plates (Sigma C7521) with 2 ml of TS medium [DMEM/F12 (Invitrogen 31330) supplemented with 0.1m 2-mercaptoethanol (Gibco 31350), 0.2% FBS (ThermoFisher Scientific 10270106), 100µg/ml Primocin (Invivogen ant-pm1), 0.3% BSA (Sigma A8412), 1% ITS-X supplement (Gibco 51500-056), 1.5 mg/ml L-ascorbic acid (Sigma A4403), 50 ng/ml EGF (Peprotech AF-100-15), 2 mM CHIR99021 (R&D 4423), 0.5 mM A83-01 (Stem Cell Tech. 72024), 1 mM SB431542 (Tocris 1614), 0.8 mM VPA (Sigma P4543) and 5 mM Y27632 (Stem Cell Technologies 72304). The medium was changed every two days, and cells were dissociated with TrypLE (Gibco 12604-021) for 10-15 min at 37°C to passage them. EVT differentiation was achieved through a modification of a protocol described previously (Okae et al. 2018). hTSC were cultured in pre-coated 6-well plates (1 µg/ml Col IV) with 2 ml of EVT differentiation medium (EVTM: DMEM/F12, 0.1 mM 2-mercaptoethanol (Gibco 31350)), 100 µg/ml Primocin (Invivogen ant-pm1), 0.3% BSA (Sigma A8412), 1% ITS-X supplement (Gibco 51500-056), 2.5 µM Y27632 (Stemcell Technologies 72304), 100 ng/ml NRG1 (Cell Signaling 5218SC), 7.5 µM A83-01 (Tocris Biotechne 2939) and 4% knockout serum replacement (KSR) (ThermoFisher 10828010). Matrigel (Corning 356231) at 2% final concentration was added as cells were suspended in medium and seeded in the plate. After 3 days, EVTM was changed and replaced with new EVTM without NRG1 and a final Matrigel concentration of 0.5%. At 6 days of differentiation, EVTM was replaced with new EVTM without NRG1 and KSR. EVTs were cultured two more days and then collected for RNA and protein extraction.

### Human samples

The placental samples from normal first and early second trimester, and normal term pregnancies used for this study were obtained with written informed consent from all participants in accordance with the guidelines in The Declaration of Helsinki 2000. Elective terminations of normal pregnancies were performed at Addenbrooke’s Hospital under ethical approval from the Cambridge Local Research Ethics Committee (04/Q0108/23). Samples were either snap-frozen for RNA isolation or embedded in formalin-fixed paraffin wax for tissue sections (4µm).

### Lentiviral transduction

For the production of lentiviral particles, 10^6^ HEK293T cells seeded in 100mm plates were cotransfected (TransIT, Mirus BIO 2700) with 6.5 µg of psPAX2 (Addgene 12260), 3.5 µg of pMD2.G (Addgene 12259), and 10µg of the appropriate lentiviral vector: dCas9-VP64_Blast (Addgene 61425), MS2-p65-HSF1_Hygro (Addgene 61426), sgRNA(MS2)_puro (Addgene 73795) cloned with an individual sgRNA. 48h later, 10 mL of virus supernatant was filtered through a 0.45 um filter (Sartorius, 16533) and supplemented with 8µg/ml polybrene (Millipore, TR-1003-G).

### Western blot

Cells were lysed in radioimmunoprecipitation assay (RIPA) buffer (20 mM Tris-HCl, pH 8.0, 137 mM NaCl, 1 mM MgCl2, 1 mM CaCl2, 10% glycerol, 1% NP-40, 0.5% sodium deoxycholate, 0.1% sodium dodecyl sulphate), containing a protease inhibitor cocktail (Sigma P2714), and incubated at 4°C for 1 h, followed by centrifugation (9300 × g, 10 min). Western blotting was performed as described previously (Perez-Garcia et al. 2014). Blots were probe against the antibodies anti-Bap1 (Cell signalling, D1W9B #13187), anti-beta-actin (Abcam ab6276), anti-tubulin (Abcam ab6160), anti-EZH2 (Diagenode, pAB-039-050), Anti-SUZ12 (NEB, 3737), Anti-ASXl1 (Cell Signaling, D1B6V #52519), anti-ASXl2 (Abcam ab106540) and anti HLA-G (Bio-rad MCA2043) followed by horseradish peroxidise-conjugated secondary antibodies anti-rabbit (Bio-Rad 170–6515), anti-mouse (Bio-Rad 170–6516), all 1:3,000). Detection was carried out with enhanced chemiluminescence reaction (GE Healthcare RPN2209) on X-ray films. The intensity of the bands was quantified using ImageJ software.

### Immunohistochemistry

Immunohistochemistry on sections of E9.5 wild-type and *Bap1* KO placentas from the DMDD collection (dmdd.org.uk) and first trimester placentas was performed as in (Turco et al. 2018). Briefly, immunohistochemistry was carried out using heat-induced epitope retrieval buffers (A. Menarini) and Vectastain avidin-biotin-HRP reagents (Vector Lab PK-6100). Anti-BAP1 antibody (Cell signalling, D1W9B #13187) was used at 1:200. For each experiment, a negative control was included in which the antibody was replaced with equivalent concentrations of isotype-matched rabbit IgG. Images were taken with an EVOS M5000 microscope (Thermo Fisher Scientific).

### Immunofluorescence staining

Cells were fixed with 4% paraformaldehyde (PFA) in PBS for 10 min and permeabilized with PBS, 0.3% Triton X-100 for 10 min. Blocking was carried out with PBS, 0.1% Tween 20, 1% BSA (PBT/BSA) for 30 min, followed by antibody incubation for 60 min. Primary antibodies and dilutions (in PBT/BSA) were: E-Cadherin (CDH1) 1:200 (BD Biosciences, 610181), anti-BAP1 1:200 (Cell Signaling, D1W9B #13187), anti-ESRRB 1:200 (R&D Systems H6707), and Phalloidin 1:500 (Thermo Scientific A12380).

Whole-mount embryo staining was performed following a modification of the protocol previously described (Kalkan et al. 2019). Briefly, dissected E6.5 conceptuses were fixed for 1 hour in 4% PFA. After three washes (15 min) with PBS supplemented with 3 mg/ml polyvinylpyrrolidone (PVP) (Sigma, P0930), embryos were permeabilized with PBS containing 5% DMSO, 0.5% Triton X-100; 0.1% BSA; 0.01% Tween 20 for 1 hour. Then, embryos were blocked overnight at 4°C in permeabilization buffer, containing 2% donkey serum. Embryos were incubated overnight at 4°C with antibodies against E-Cadherin (CDH1) at 1:200 (BD Biosciences, 610181) and BAP1 at 1:100 (Cell Signaling, D1W9B #13187) in blocking buffer, followed by three washes in blocking buffer for 1 hour. Then, conceptuses were incubated overnight with secondary Alexa Fluor 488 or 568 (Thermo Fisher Scientific) antibodies diluted 1:400 in blocking buffer. Lastly, embryos were washed three times for 1 hour in blocking buffer and nuclei were counter-stained with DAPI. For embryo mounting, samples were taken through a series of 25%, 50%, 75% and 100% Vectashield (Vector Laboratories, H-1000) diluted in PBS. Embryos were mounted in Vectashield, surrounded by spacer drops of vaseline for the coverslip, to immobilise embryos.

Images were taken with an Olympus BX61 epifluorescence microscope or a Zeiss LSM 780 confocal microscope. Images were processed with Fiji software.

### Cell Adhesion assay

Adhesion capacity of vector-control and *Bap1*^-/-^ mTSCs was measured using the Vybrant cell adhesion assay kit (Thermo Fisher Scientific V13181) as previously described (Branco et al. 2016). Briefly, cells resuspended in serum-free RPMI medium were labelled with Calcein AM (5uM) during 30 min at 37°C. Cells were washed twice with RPMI medium and 10^5^ cells plated per well in a 96-well tissue culture plate and left to attach for 2 hr in serum-free RPMI medium. Finally, cells were washed three times and the remaining attached cells were detected measuring the fluorescence emission at 517 nm with a PHERAstar FS plate reader.

### Trophosphere generation

Trophosphere were generated following a modification of a protocol described previously (Rai and Cross 2015a). In brief, 10^4^ WT and mutant cells resuspended in complete medium were cultured in Ultra-Low Attachment plates (Corning, USA). 48h later, cells were collected, washed with PBS and transferred back to Ultra-Low Attachment dishes with differentiation medium for another 7 days. Then, the trophospheres were collected for RNA analysis.

### Trophoblast cell invasion

The invasion assays were carried out following a modification of the protocol described in (Hemberger, Hughes, and Cross 2004). The Transwell filters (Sigma, CLS3422) were coated with 100 µl of a 1:20 dilution of cold Matrigel (Corning 356231) in RPMI 1640 medium. The Matrigel layer was allowed to dry overnight at room temperature and was rehydrated the next day with 100 µl of supplemented RPMI 1640 medium for 2 h at 37°C under 95% humidity and 5% CO2. Confluent 60-mm dishes of TS cells were trypsinized and resuspended in RPMI at 10^6^ cells/ml. 100 µl of this cell suspension (10^5^ cells) was added to the top chamber, and the bottom chamber was filled with 800 µl of culture medium.

After the specific times of incubation, Transwell inserts were fixed for 5 min in 4% paraformaldehyde and washed with 1× PBS. Cells that remained on top of the filters as well as the Matrigel coating were scraped off. Filters were stained overnight with hematoxylin and excised under a dissecting microscope, removing all residual cells from the top of the filters. Filters were mounted with 20% glycerol in 1× PBS and photographs of each filter were quantified with Image J.

### Proliferation assay

Analysis of cell proliferation rate was performed as in (Woods et al. 2017). In brief, 10,000 vector-control and *Bap1* KO mTSCs were plated in complete medium and collected every 24 h over 4 days. After trypsinisation, the number of viable cells was counted using the Muse Count & Viability Assay Kit (Merck Millipore MCH100102) and run on the Muse cell analyser (Merck Millipore), according to manufacturer’s instructions. Statistical analysis was performed using ANOVA followed by Holm-Sidak’s posthoc test.

### RT-qPCR

Total RNA was extracted using TRI reagent (Sigma T9424), DNase-treated and 1µg used for cDNA synthesis with RevertAid H-Minus reverse transcriptase (Thermo Scientific EP0451). Quantitative (q)PCR was performed using SYBR Green Jump Start Taq Ready Mix (Sigma S4438) and Intron-spanning primer pairs (Supplementary Table 5) on a Bio-Rad CFX384 thermocycler. Normalised expression levels are displayed as mean relative to the vector control sample; error bars indicate standard error of the means (S.E.M.) of at least three replicates. Where appropriate, Student’s t-tests or ANOVA were performed to calculate statistical significance of expression differences (*p* < 0.05) using GraphPad Prism 7.

### RNA-seq

For RNA-seq, total RNA was extracted with Trizol followed by DNase treatment using TURBO DNA-free kit (Life Technologies AM1907). For wild-type and *Bap1* KO mTSC experiments, adapter indexed strand-specific RNA-seq libraries were generated from 1000 ng of total RNA following the dUTP method using the stranded mRNA LT sample kit (Illumina). Libraries were pooled and sequenced on Illumina HiSeq2500 in 75bp paired-end mode. FASTQ files were aligned to the Mus musculus GRCm38 genome reference genome using HISAT2 v2.1.0. Sequence data were deposited in ENA under accession ERP023265.

For RNA-seq from SAM *Bap1*-overexpressing cells, RNA-seq libraries were generated from 500 ng using TruSeq Stranded mRNA library prep (Illumina, 20020594). Indexed libraries were pooled and sequenced on an Illumina HiSeq2500 sequencer in 100 bp single-end mode. FastQ data were map to Mus musculus GRCm38 genome assembly using HISAT2 v2.1.0.

### Bioinformatic analysis

Data were quantified using the RNA-seq quantitation pipeline in SeqMonk (www.bioinformatics.babraham.ac.uk), and normalised according to total read count (read per million mapped reads, RPM). Differential expression was calculated using DESeq2 and FPKM Fold Change ≥2 with P<0.05 and adjusted for multiple testing correction using the Benjamini– Hochberg method. Stringent differential expression was calculated combining DESeq2 and intensity difference filters in SeqMonk. Bap1 KO cell lines, data were retrieved from ENA accession number ERP023265 and (He et al. 2019).

Heatmaps and Principal component analysis were generated using Seqmonk. Gene Ontology (GO) was performed on genes found to be significantly up-or downregulated, against a background list of genes consisting of those with more than ten reads aligned. GO terms with a Bonferroni P-value of <0.05 were found using DAVID (Dennis et al. 2003). Venn diagrams were plotted using BioVenn (www.cmbi.ru.nl/cdd/biovenn/). The significance of the pairwise overlap between the datasets was determined by the overlapping gene group tool provided at www.nemates.org.

S

### Statistics

### Data availability

Genome-wide sequencing data are deposited in the GEO database under accession number (to be confirmed); they are also available directly from the authors.

## Acknowledgments

We would like to thank Dr Anne Segonds-Pichon for expert help with statistical analyses, the Flow Cytometry Facility at the Babraham Institute and Kristina Tabbada for Illumina high-throughput sequencing.

This work was supported by the Wellcome Trust Strategic Award WT100160MA and by the Centre for Trophoblast Research, University of Cambridge, UK. V.P.-G is the recipient of a Next-Generation Fellowship awarded by the Centre for Trophoblast Research, University of Cambridge, UK. P.L.-J is supported through an Erasmus+ traineeship.

## Competing interests

The authors have no competing interests to declare.

## Author contributions

V.P.-G and P.L.-J performed the bulk of the experiments and analysed the data; G.J.B., A.M. and M.Y.T. provided intellectual input and, reviewed and edited the manuscript. V.P.-G and M.H. designed experiments and wrote the manuscript with input from all the authors.

---

**Figure 1-figure supplement 1:**
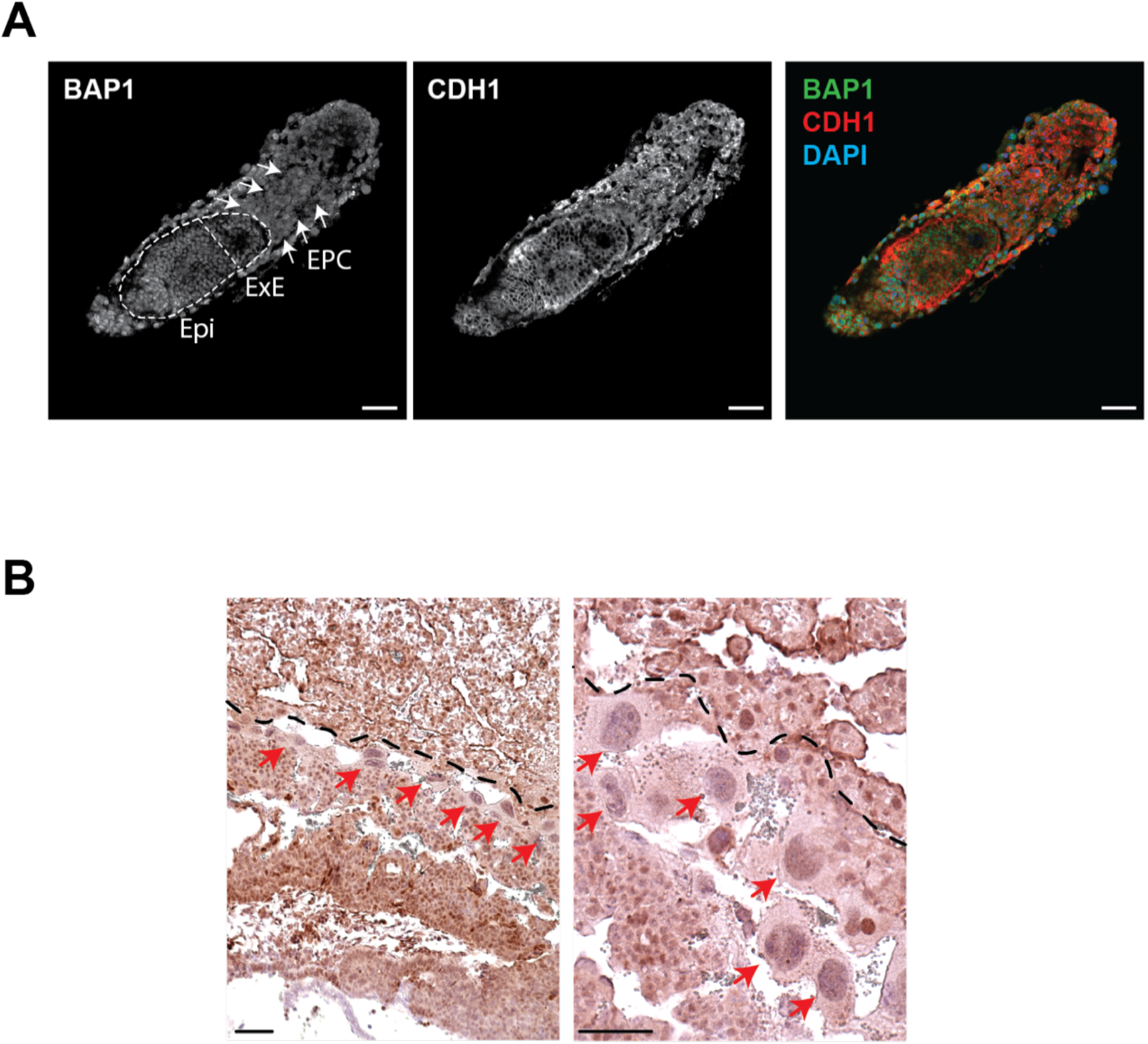
Bap1 expression in early mouse placentation. A) Whole-mount immunostaining of E6.5 conceptuses for BAP1 and CDH1. Bap1 is highly expressed in the Epiblast (Epi) and extraembryonic ectoderm (ExE). The ectoplacental cone (EPC) shows a reduced and diffuse staining (highlighted by arrows) as differentiation progresses. Scale bar: 100 µm. B) Immunohistochemistry analysis of E9.5 placenta for BAP1. Red arrows highlight trophoblast giant cells. The dotted lines separate the decidual tissue (upper part) from the rest of the placenta. Scale bar, 100 µm.

**Figure 2-figure supplement 1:**
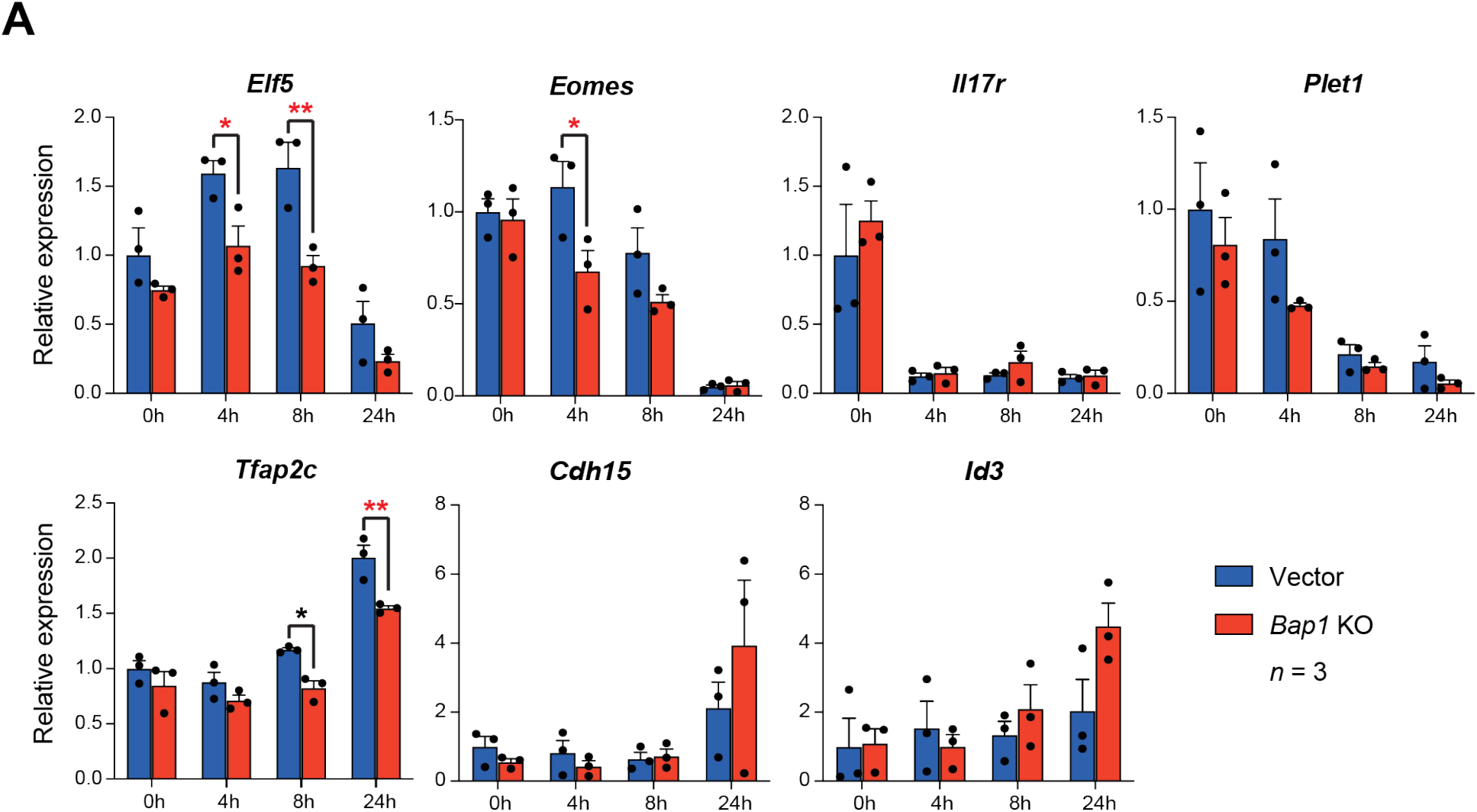
Bap1 ablation does not negatively affect stemness. A) RT-qPCR analysis to assess the effect of Bap1 ablation on the stem cell state and early differentiation of mTSCs. Data are normalised to Sdha and are displayed as mean of three biological (=independent clones) replicates ± S.E.M.; *p < 0.05, **p < 0.01 (two-way ANOVA with Sidak’s multiple comparisons test).

**Figure 3-figure supplement 1:**
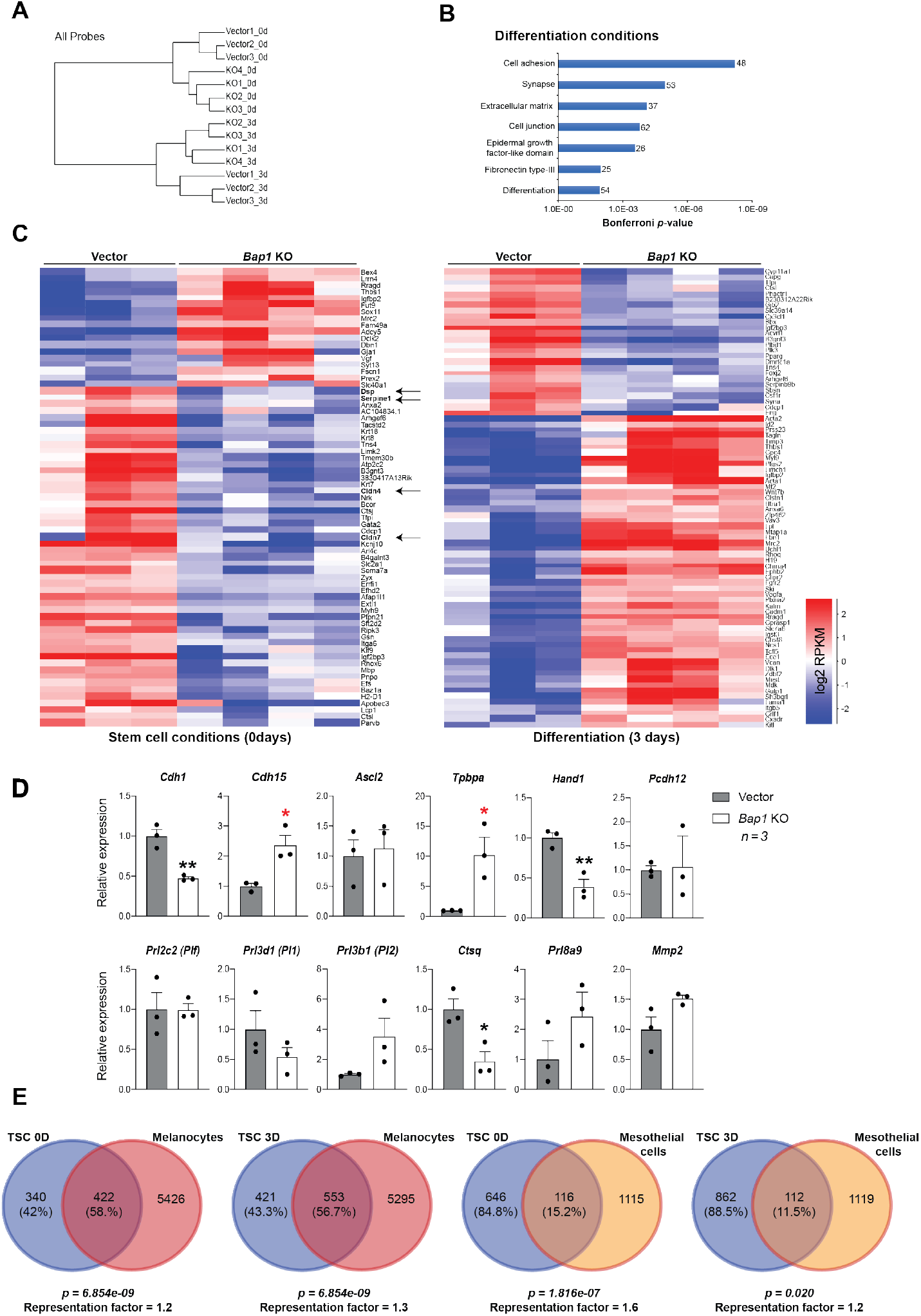
Bap1-deficicency induces precocious EMT with features of malignant transformation. A) Hierarchical clustering of global transcriptomes generated by RNA-seq from 3 independent vector-control and 4 Bap1 knockout (KO) mTSC clones grown in stem cell conditions (0d) and after three days of differentiation (3d). B) Gene ontology analyses of genes differentially expressed between 3-day differentiated vector-control and Bap1-mutant mTSCs. C) Heatmap of differentially expressed genes as in (A). Arrows highlight de-regulated genes essential for the stabilization of cell-cell contacts and epithelial integrity. D) RT-qPCR analysis of 3D-trophospheres generated from vector-control and Bap1-mutant mTSCs after 8 days of differentiation. Data are normalised to Sdha and are displayed as mean of three replicates ± S.E.M.; * p < 0.05, ** p < 0.01 (Student’s t-test). E) Venn diagrams of genes de-regulated in common in Bap1-mutant mTSCs (0d and 3d) and Bap1-null melanocytes and mesothelial cells (He et al. 2019).

**Figure 4-figure supplement 1:**
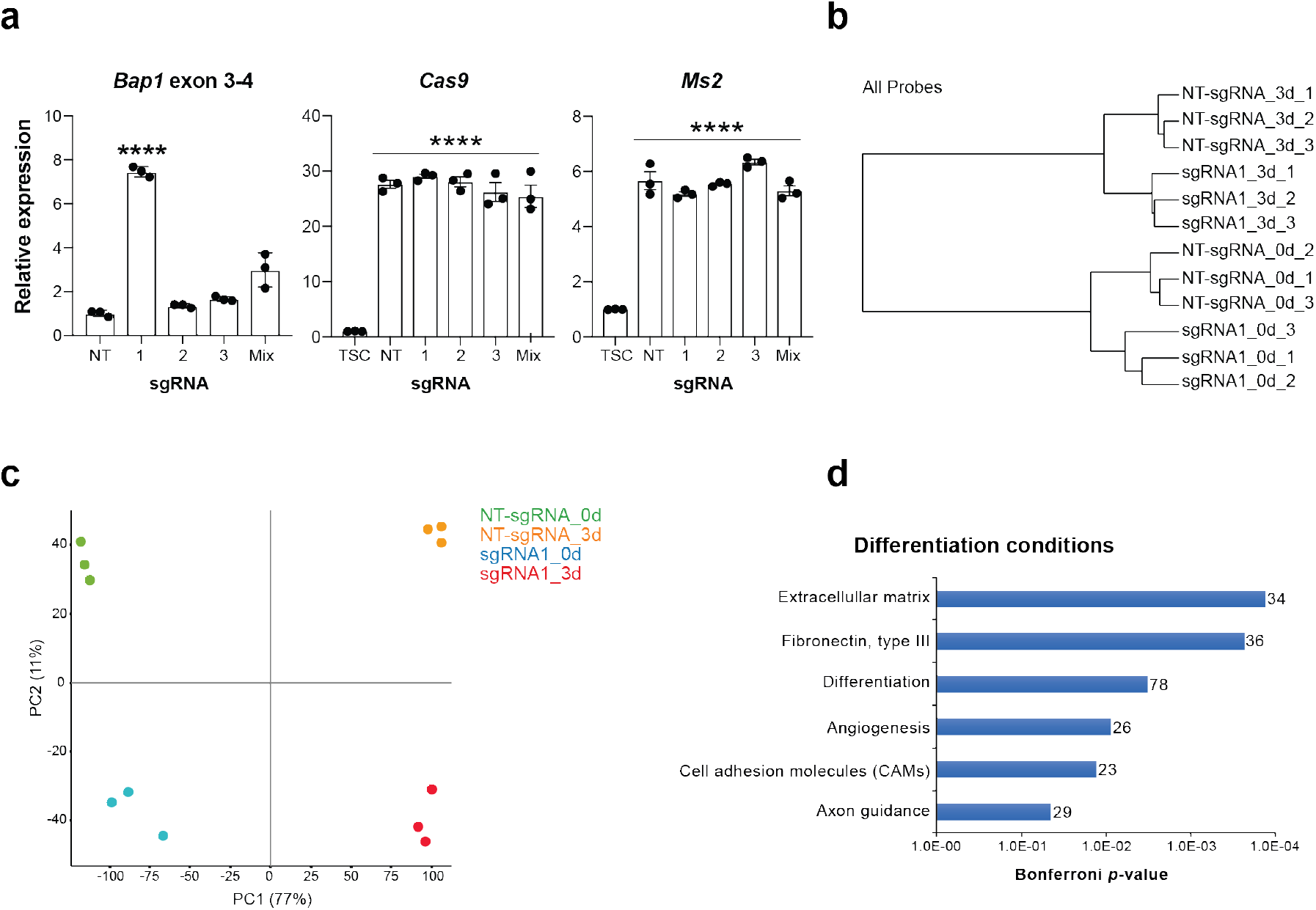
Bap1 overexpression increases epithelial features of mTSCs. A) First graph shows RT-qPCR analysis to determine the overexpression of Bap1 (exon 3-4) in mTSCs induced by the transduction of each single guide RNAs (sgRNAs) or in combination (Mix) compared to non-targeting sgRNA (NT). Second and third graphs show the stable overexpression of the Cas9 and Ms2 SAM components for each cell line generated compared to non-transduced mTSCs examined by RT-qPCR. Data are normalised to Sdha and are displayed as mean of three replicates ± S.E.M.; ****p < 0.0001 (one way-ANOVA with Dunnett’s multiple comparisons test). B) Hierarchical clustering analysis of global transcriptomes generated by RNA-seq from NT-gRNA and sgRNA1 mTSCs grown in stem cells conditions (0d) and after three days of differentiation (3d). Samples cluster according to the amount of BAP1 and day of differentiation. Three independent replicates were sequenced in each condition. C) Principal component analysis of global transcriptomes as in (B). D) Gene ontology analyses of genes differentially expressed between sgRNA1- and NT-sgRNA mTSCs differentiated for 3 days.

**Figure 5-figure supplement 1:**
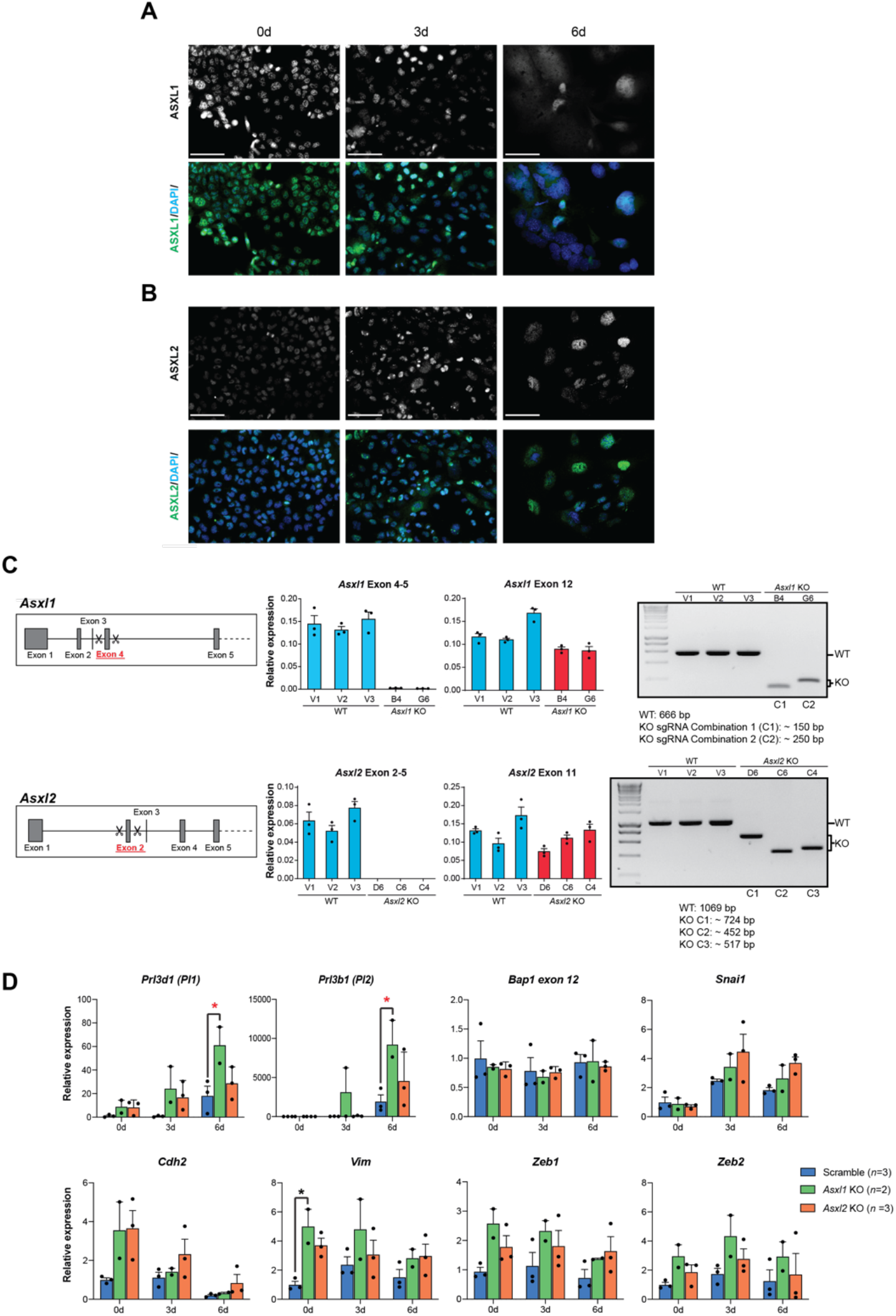
CRISPR-mediated KO of Asxl1 and Asxl2 in mTSCs. A) Immunofluorescence analysis for ASXL1 on mTSCs in stem cell conditions and differentiated for 3 and 6 days. Data confirm that ASXL1 is downregulated during trophoblast differentiation. Scale bars: 100 µm. B) ASXL2 immunostaining as in (A) shows that the levels of ASXL2 increase as mTSCs differentiate. Scale bars: 100 µm. C) Details of the CRISPR/Cas9 knockout strategy for ablating Asxl1 and Asxl2 genes. RT–qPCR and genomic genotyping PCR analysis were performed on single-cell expanded mTSCs clones to confirm homozygous knockout. Data are mean ± S.E.M. of n = 3 technical replicates. D) Additional RT-qPCR analysis of Asxl1 and Asxl2 KO mTSCs clones as in Figure 5. Data are mean ± S.E.M. of n = 3 (wild-type, scramble), n = 2 (Asxl1 KO) and n = 3 (Asxl2 KO) individual clones as independent replicates; *p < 0.05 (two-way ANOVA with Sidak’s multiple comparisons test).

**Figure 6-figure supplement 1:**
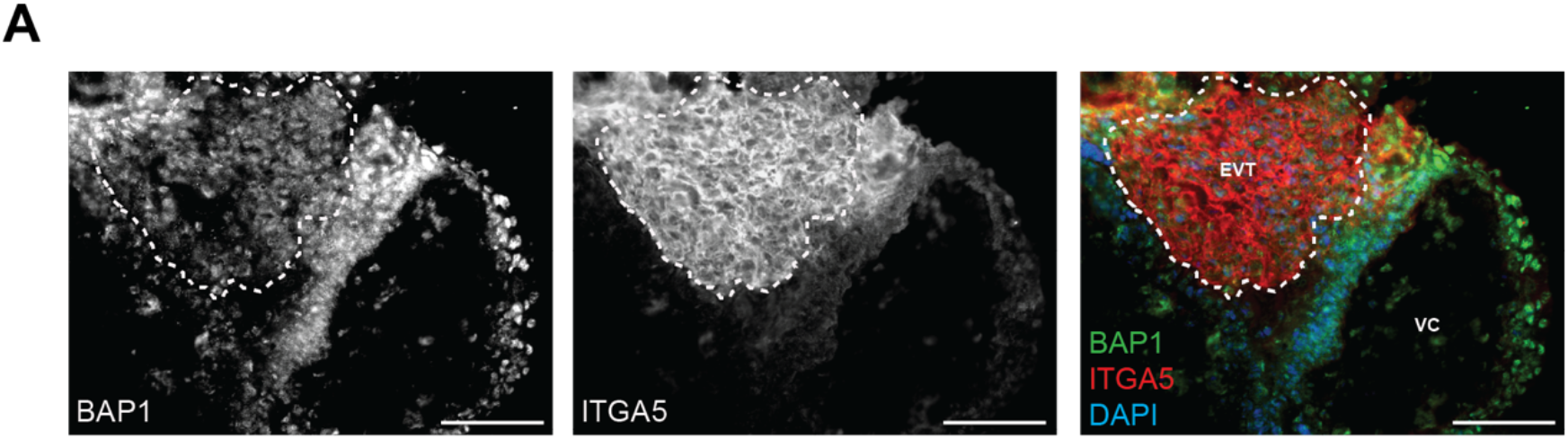
BAP1 immunofluorescence staining of first trimester human placenta. A) Panels show staining of BAP1 combined with Integrin alpha-5 (ITGA5). Dotted lines demarcate the area of extravillous trophoblast (EVT) demarcated by high expression of ITGA5. VC: villous core. Scale bar: 100 µm.

**Supplementary Table 1: Genes dysregulated in Bap1 KO mTSC in stem cell conditions (0d) and during differentiation (3d).** Excel file uploaded as a Supplementary file.

**Supplementary Table 2: Gene ontology analyses of genes differentially expressed between vector and *Bap1*-mutant mTSCs in stem cell conditions (0d) and during differentiation (3d).** Excel file uploaded as a Supplementary file.

**Supplementary Table 3: Genes dysregulated in Bap1 overexpressing mTSC in stem cell conditions (0d) and during differentiation (3d).** Excel file uploaded as a Supplementary file.

**Supplementary Table 4: Gene ontology analyses of genes differentially expressed between Bap1 overexpressing and control mTSCs in stem cell conditions (0d) and during differentiation (3d).** Excel file uploaded as a Supplementary file.

**Supplementary Table 5:**
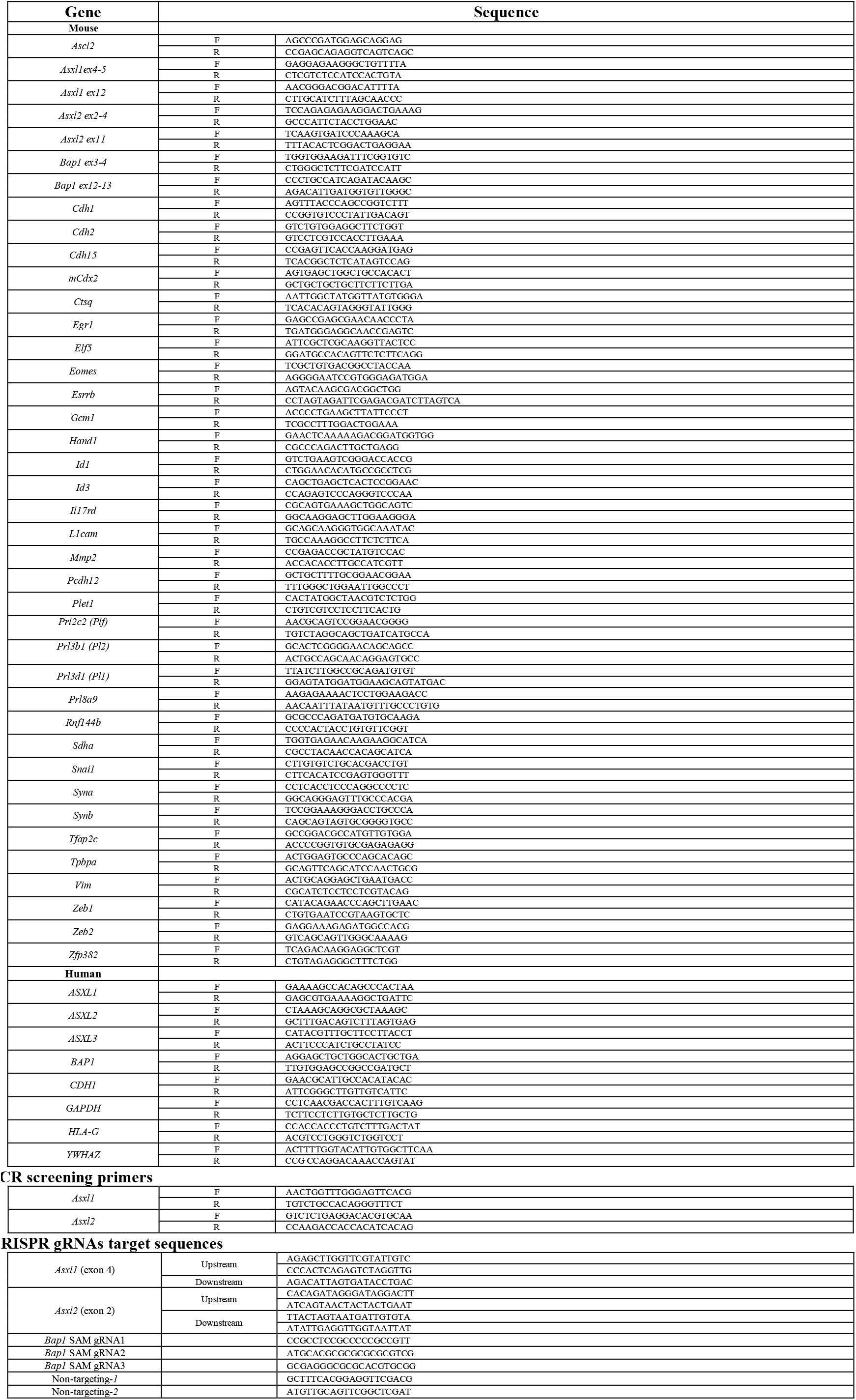
Primer sequences for RT-qPCR and CRISPR gRNAs target sequences.

